# A high-throughput method for measuring fungal growth rate on solid media using automated imaging and deep learning

**DOI:** 10.64898/2026.03.04.709619

**Authors:** Thea Kristensen, Erik Bjørnager Dam, Henrik H. De Fine Licht

**Author notes:** **Author for correspondence**:, Address: Section for Organismal Biology, Department of Plant and Environmental Science, University of Copenhagen, Thorvaldsensvej 40, 1871 Frederiksberg, Denmark.

## Abstract

Measuring the growth rate of filamentous fungi is an essential phenotype assay in fungal biology, enabling the comparison of nutrient-related fitness metrics across various isolates, species and genera. Conventional methods are time consuming and labor intensive, which prohibits the adaptation and implementation of high-throughput phenotyping. Here, we suggest a high-throughput methodological pipeline to study fungal growth on solid media combining the use of 24-well plates, an automated image acquisition system, and human assisted deep learning analysis of acquired images. Training a deep learning model through an iterative process – with continuous feedback and corrective annotations – enabled the development of a satisfying model that automatically segments pixels belonging to either fungus or background within a few hours. We evaluated this deep learning model by applying it to two test sets: First, a set of 336 images was used to validate the results by comparison with manual measurements. We demonstrate that the automated segmentation approach provides robust estimation of fungal growth not significantly different to manually segmented data. Second, a larger test set consisting of 2,016 images was used to illustrate the scalability of the model. After training the model for less than two hours, the deep learning model segmented the entire image data set automatically within minutes. The presented method is easily scalable and adjustable to other fungi and growth morphologies, due to the interactive training. Moreover, by combining 24-well plates and automatic image acquisition, measurements can be sped up as growth is detected across a smaller surface area than a standard six or nine cm diameter petri dish. The proposed methodological pipeline thus offers a new tool for estimating fungal growth rates, which can accelerate measurements, reduce bias, and increase throughput.

## Introduction

The fungal kingdom consists of a considerable diversity of species, occupying many different ecological niches and taking on a myriad of ecological roles (Blackwell, 2011). From important decomposers, over highly optimized production strains in the biotechnology industry to detrimental pathogens of humans, animals and plants (Naranjo-Ortiz & Gabaldón 2019). With genetic data becoming cheaper and faster, an increasing amount of data is available to understand more about these fascinating organisms and a new bottleneck is to gather enough phenotypic data to match the genetic data (Anderson and Kohn 1998, Milgroom 2017). A widely used phenotyping assay in mycology is to measure how fast a given isolate grows in liquid or on agar media, as this provides a nutrient related fitness metric that can be compared between species and genera (Anderson et al. 2019, Pringle and Taylor 2002, Slowik et al. 2023, Trinci 1971). The growth media can vary in terms of concentration of nutrients, antibiotics, pH levels, or the growth assay can be executed at varying temperatures, CO_2_ levels and/or humidity to identify context-dependent differences in growth rate.

The conventional method to obtain a fungal growth rate on agar media is to use a centrally placed fungal inoculum in a petri dish containing agar and manually measure growth in terms of diameter or radius of the growing mycelium. This can be measured with a ruler or by sequential photographs that can be digitally analysed, for example using the widely used software ImageJ (Cánovas et al. 2017, Lendenmann et al. 2014, Schneider et al. 2012, You and Chung 2007). This method is straightforward and allows for other phenotypic characters such as colour or sporulation to be recorded, while differences in mycelial density is generally not captured. Manually measuring fungal colony size directly from plates or from pictures is a labour intensive and time-consuming task, why the change in growth is usually only recorded once per day or as an end point several days after inoculation. Systems to automatically monitor fungal growth over time exist, mainly based on growth in liquid media using e.g., optical density (OD) or microscopically tracking hyphal extension (Granade et al. 1985, Lee et al. 2021, Slowik et al. 2023). Studying submerged cultures may be an artificial reflection of their phenotype, as filamentous fungi naturally grow on or in many different solid or composite substrates. Moreover, by measuring growth in liquid cultures, important information (such as colour, morphology, sporulation, and potential inhibition zones in co-cultures) will often be missed.

Automated imaging is increasingly used for data acquisition and storage as it enables the capture of time-specific data at all hours of the day, thus enabling detailed data in high time resolution. Manual annotation of such time-series image data quickly becomes unfeasible as time spent on measuring fungal growth scales linearly with number of isolates, time steps, and plates or wells studied. The combination of automated image acquisition and segmentation further allows for more rapid growth measurements to be obtained, as images with just a few hours in between can be measured and analyzed. This is often not feasible with manual measurements except for very rapidly growing fungi (Slowik et al. 2023). There is thus a growing need for computer-aided analysis to ultimately increase throughput of fungal growth phenotyping, by reducing length of experiments and time spent on measurement, from several days and even weeks to 1-2 days.

Here, we show that a semi-automatic, high-throughput methodological pipeline to study fungal growth on solid media provides robust estimation of fungal growth comparable to manually measured data. The method combines an automated image acquisition system with freely available deep learning software called *RootPainter* (Smith et al. 2022). We generated fungal growth data consisting of time-series images of isolates from the plant pathogenic fungal genus *Colletotrichum* on solid agar media. Any automatic or manual image acquisition system can be used, while in this specific set-up we used a Reshape Imaging System (RIS) 1.0 (Reshape Biotech Inc., Copenhagen). The image data was processed to define patterns of filamentous fungal growth using semi-automatic deep learning segmentation. In just a few hours of human-assisted training we generated a deep learning model that can successfully detect fungus-covered area on full datasets of growth-experiment images. The areas detected by the model were further processed in *R* (R Core Team 2022) to automatically fit growth models and extract the maximum growth rate. We compare the applicability of this methodological pipeline to manual measurement of fungal growth image data and demonstrate how the presented method provides growth data of similar quality.

## Materials and Methods

### Fungal isolates and generation of conidia suspensions

Four isolates of the plant pathogenic fungal genus *Colletotrichum* were used in the experiments: *C. gloerosporiodes s. lat*. (ESALQ-C005), *C. gloerosporiodes s. lat*. (ESALQ-C006), *C. acutatum s. lat*. (ESALQ-C008), and *C. acutatum s. lat*. (ESALQ-C009). Isolates were revived from −80 °C glycerol stocks and grown on standard petri dishes (diameter: 90 mm) containing Potato Dextrose Agar (PDA) medium composed of 39 g Potato Glucose Agar (Sigma-Aldrich) (potato extract 4g, dextrose 20 g, agar 15 g) in 1 L of distilled water. Conidia suspensions were obtained by inoculating a new 90 mm diameter PDA plate and growing fungal cultures for five days at 25°C. Conidia were harvested by covering the plate with 10 mL of 0.05 % Triton X and gently loosening spores with a drigalski spatula. This suspension was transferred to a 50 mL centrifuge tube, which was centrifuged at 3000 RPM for 3 minutes at 5° C. The supernatant was discarded, and the spores were washed by adding 10 mL 0.05 % Triton X, vortexing at low power, and centrifugation at 3000 RPM for 3 minutes at 5° C and discarding the supernatant. Washing was repeated once with 10 mL 0.05 % Triton X and finally with 10 mL demineralized water. Subsequently, the spores were resuspended in 2-5 mL demineralized water to obtain a stock solution. The concentration of the stock solution was determined by counting spores in parallel imaging scanning through 100 μl triplicate dilutions in a flat-bottom 96-well plate, which thereby generated series of 10 images of each well. The images with a total scan area length of 405 μm, were measured using an oCelloScope™ microscope detection system (objective, 4 ×) (Biosense Solutions Aps, Copenhagen) and the number of spores in each well were estimated using the segmentation tool in the associated software *UniExplorer* (v. 11.0.1.8356). The concentration of the stock solution was calculated from each well and the average across replicate wells used. The stock solution was then diluted to reach a concentration of 1 x 10^6^ spores/mL which was used for inoculation.

### Generating experimental fungal growth data sets

The four fungal isolates were cultivated on PDA medium with a standard pH of 5.6 and on PDA medium adjusted to pH 3. The assay was set up using 24-well plates (Greiner BIO-ONE GmbH, Kremsmünster, Austria) (individual well diameter: 16 mm), with each well containing 1.5 mL PDA medium (Fig. 1A). Each well was inoculated with 1 μL of spore suspension with pre-set concentrations of 1 x 10^6^ spores/mL (corresponding to 1,000 conidia inoculated per well). Each isolate was inoculated in six wells per plate, corresponding to six technical replicates for each growth media type. Images were obtained using a Reshape Imaging System (RIS) 1.0 (Reshape Biotech Inc., Copenhagen), which has a mobile arm with a camera and light source attached that moves over the plates and takes pictures of the individual wells. The RIS was set to acquire a picture of each well every second hour for 84 hours in total with ‘2 x top light’ using a black background. This resulted in 42 images per well and image specifications are 2,000 x 2,000 pixels, RGB color encoded, JPEG format. The RIS was placed in an incubator set to 25°C and the temperature was measured throughout the experiment. A total of 2,016 images were generated in the experiment, and these were divided in three datasets: a training set, and two test sets. The Training set consisted of 1/8 of all images generated in the experiment (amounting to 256 images in total), selected as every eighth timepoint per well. Test set 1 consists of all images of a single replicate from the four isolates on both pH levels (amounting to 8 x 42 = 336 images), and Test set 2 consists of all 2,016 images. Images in the training set overlap with both test sets (Fig. 1B).

**Figure 1.**
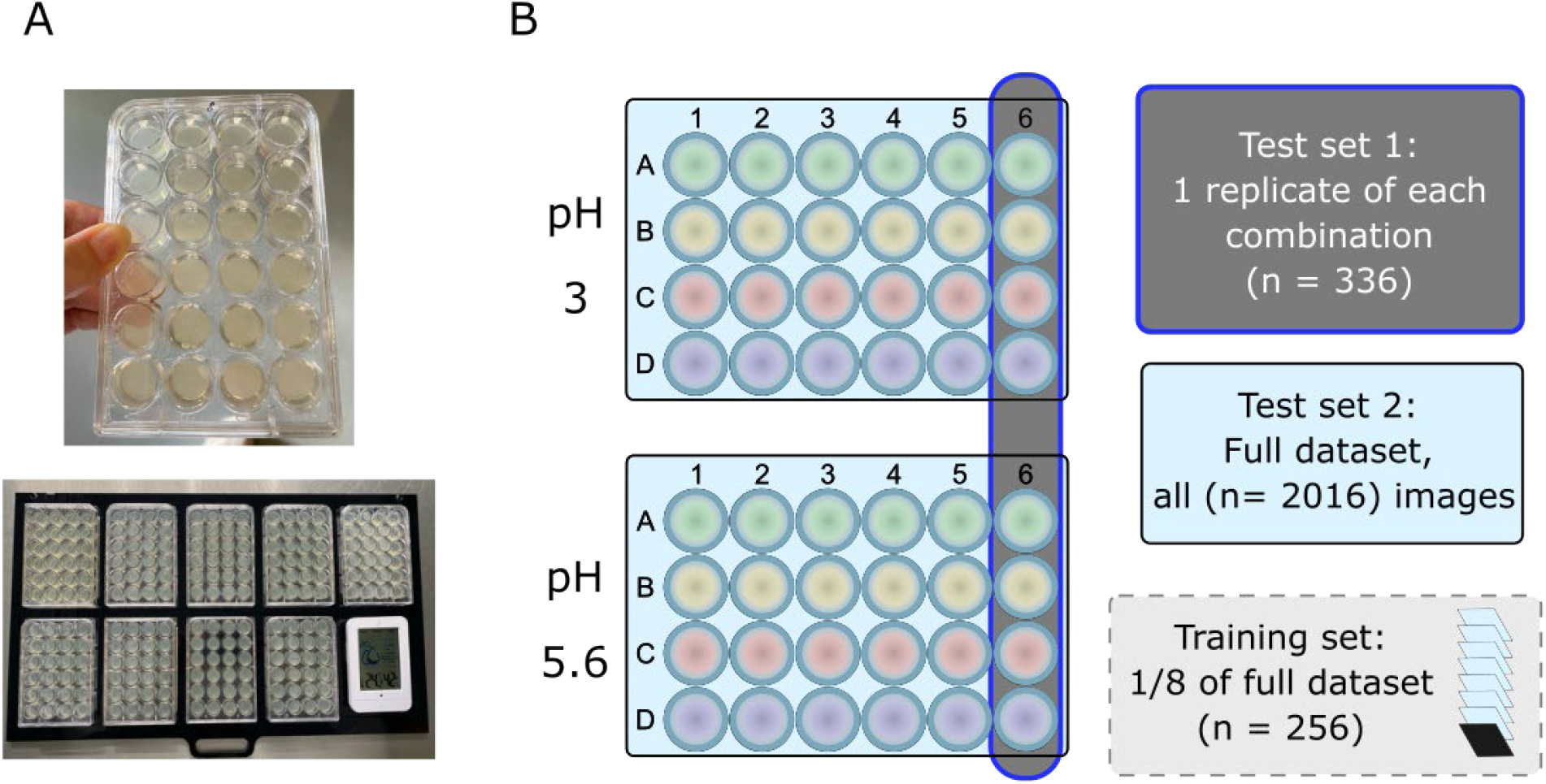
Overview of assay setup. A) Illustration of well plates prepared for imaging. B) Illustration of the experimental setup and datasets. Four isolates (A, B, C, and D) were grown on PDA medium, in six technical replicates on two pH levels (pH 3 and pH 5.6). Images were divided into the Training set consisting of 256 images (selected as every eight timepoint per well). Test set 1 consisting of 336 images and Test set 2 consisting of all 2,016 images. There is an overlap between the training set and the test sets.

### Description of the analysis pipeline (building a deep-learning model)

To automate the segmentation process of the acquired image data, a deep learning approach was used by training a model that can automatically detect which pixels in the images can be classified as fungal mycelium. We used the freely available software *RootPainter* which was developed to make the process of training a neural network more accessible (Smith et al. 2022). The software *RootPainter* operates through an iterative process, continuously running training in the background, while the user annotates images that are then used to train the model further (Fig. 2). The model is continuously improved and applied, allowing users to correct annotations instead of annotating all images from scratch. This approach saves a significant amount of time and enables observing the model’s improvement throughout the model training (Smith et al. 2022).

**Figure 2.**
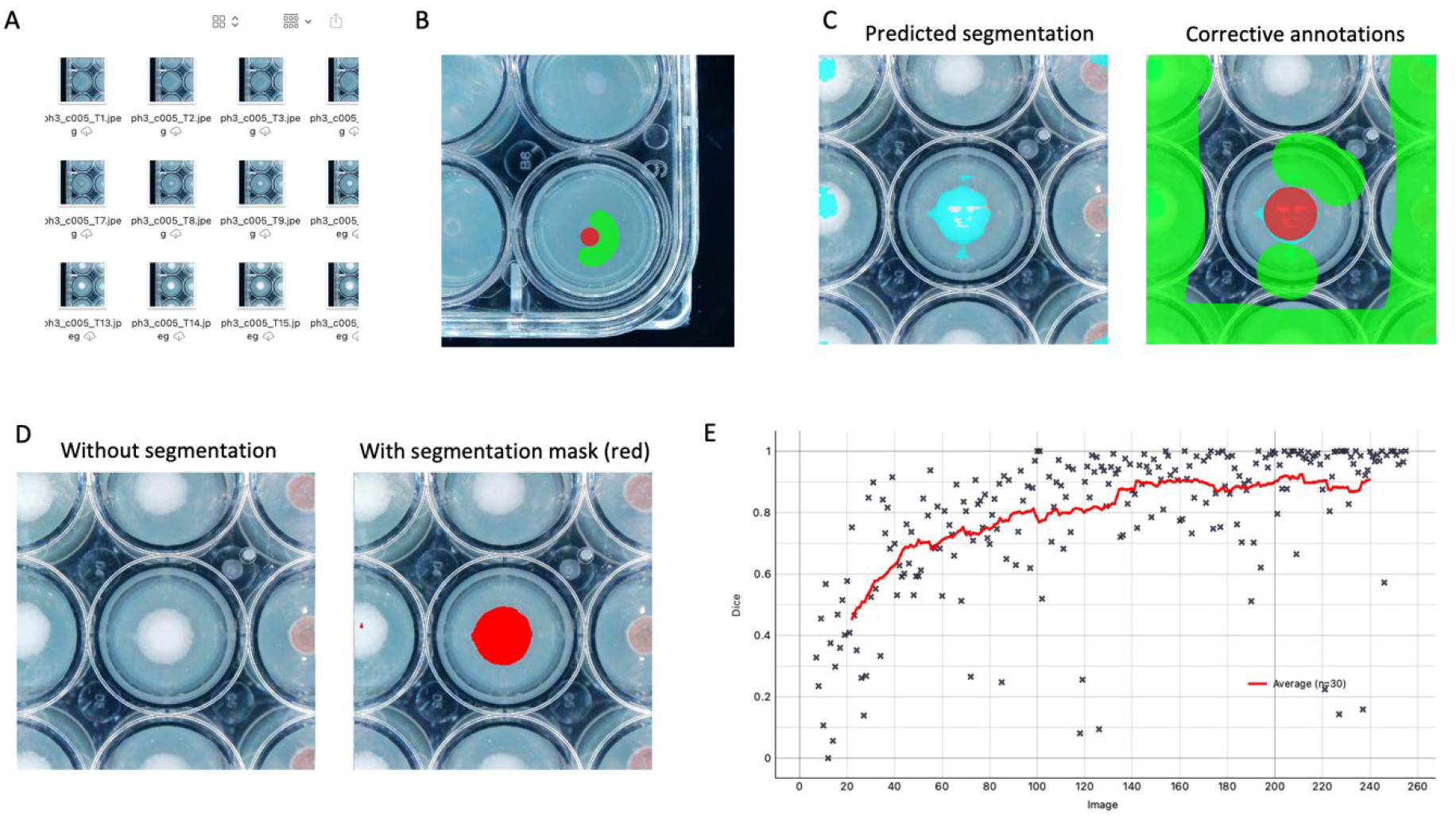
A) Directory with image files that are part of the training set. B) Rough annotation of image 1 (well diameter = 16 mm). C) Predicted segmentation while training (left) and corrected annotation of this image (right). D) Example image segmented by the final model showing without (left) and with the predicted segmentation mask (right). E) Metrics plot illustrating the accuracy of the model prediction (Y-axis) as a Dice score ranging from 0 to 1, which is higher when the model prediction agrees with ground truth, versus the number of images annotated (X-axis).

The workflow of the entire pipeline is elaborated in detail here: First, within the software *RootPainter*, the *Training set* is specified, a new model is created and two images (randomly picked by the software) are roughly annotated by the user into foreground (i.e. part of image with fungal material) and background using the built-in brush tools (Fig. 2B). This manual image annotation step does not require any coding or specialized data processing skills by the user. Next, another image (the third image) is automatically loaded by the software for manual annotation, while the automated model training runs in the background. Images are manually annotated until a second model file appears in the directory where the output model files are saved by *RootPainter*.

When the second model file appears, corrective annotation is initiated: the segmentation suggested by the newest available model is loaded, visually evaluated and the brush tools are used to manually correct false positives and false negatives on the images. Simultaneously, the software is training in the background and continuously producing new models. The metrics plot (Fig. 2E) can be evaluated while continuing to manually correct automatically annotated images. This plot shows the convergence over time (measured in images annotated) of how well the predicted segmentations fit the corrected annotations and allows the user to make the decision of when to stop training. In the example illustrated (Fig. 2E), the curve flattens after annotating approximately 140 images and does not improve even after training on 240 images. Thus, in this case, training could have been stopped earlier without noticeably affecting the resulting model.

The final model is applied to segment all images in the datasets once a satisfying model is obtained, i.e. when no further improvements appear and the average line in the metrics plot flattens (Fig. 2E). The output is a mask of the pixels classified as fungal mycelium for each image and the area of these image masks can be extracted via *RootPainter*. These measurements (area in pixels) can then be plotted as a function of time to visualize the growth of each isolate at each pH level using *R* (v. 4.1.3) and *RStudio* (Posit team 2023). The *R*-package *growthrates* (Petzoldt 2022) was used to fit a log-linear growth model to the time-series datasets for each combination of isolate and pH level, which allowed the estimation of growth parameters such as the maximum growth rate to be extracted based on the fitted models. The function *fit_easylinear* (h=4) was used to determine the subset of four datapoints with the steepest slope of the log transformed data. The slope of that subset is equal to the maximum growth rate under the assumption of exponential growth (Hall et al. 2014).

### Comparing deep learning results with manually annotated images (Test set 1)

To evaluate the workflow, all images of *Test set 1* were segmented automatically by the model and the area manually measured using the software *ImageJ* (v. 1.54g, Schneider et al. 2012). For the manual area estimation we measured the diameter of the fungus at each image by drawing a straight line across the colony and quantifying this length with the *ImageJ* function *Measure*. This was done twice for each image, with the two measurements being perpendicular. Images taken in the beginning of the growth phase, with no visible mycelium, were skipped. Manual segmentation was stopped when the growth stagnated or the edge of the well was reached. This amounted to a total of 249 images manually measured. The average diameter and the corresponding area in pixels of the fungus were calculated under the assumption that growth is circular. This is a valid assumption under our growth conditions based on manual inspection of our raw image data. To compare the results between the automatic and manual segmentation, we plotted the log-transformed area in pixels over time for both methods using *R* (v. 4.1.3). From these plots, we identified the sequence of datapoints with the steepest slope using the *R*-package *dpseg* based on a minimum of 4 continuous datapoints and a break-point penalty of 0.001 (settings: jump = FALSE, P = 0.001, minl = 4)(Machne, 2020). We used the overlapping timepoints between manually and automatically segmented data to test whether there was a significant difference between the slopes obtained from the two methods. This was done by linear regression analysis followed by pairwise slope comparisons using the R-package *lsmeans* (Lenth, 2016). The Benjamini-Hochberg method to control for false discoveries due to multiple testing was applied (Benjamini & Hochberg 1995).

### Expansion of the model to full dataset (Test set 2)

Images in *Test set 2* were automatically segmented and the area in pixels of each segmentation was extracted and imported into *R*. We plotted the log-transformed area over time, fitted a log-linear growth model and estimated growth parameters following the method of Hall et al. (2014) using the *R*-package *growthrates*. The function *fit_easylinear* (h=4) was used to determine the subset of four datapoints with the steepest slope of the log-transformed data, which is equal to the maximum growth rate under the assumption of exponential growth. Modelling was done on data where the log-transformed area in pixels is greater than 8, to avoid arbitrary variation in segmentation during the lag growth phase. The maximum growth rate was extracted for each replicate of each combination of isolate and pH level and visualized as jitter plots using the *geom_jitter* function from the *R*-package *ggplot2* (Wickham, 2016). Variability of the data within and between isolate-pH combinations were tested with a one-way ANOVA followed by a TukeyHSD post-hoc test using *R*.

## Results

Four isolates of *Colletotrichum* fungi were grown across two pH levels, 3 and 5.6, and growth estimated by acquiring images of the growing colonies every two hours (Fig. 1). We used these data to demonstrate how to automatically measure growth using a machine-learning assisted iterative process of image segmentation each pixel into either fungus or background (Fig. 2). To validate our method, we compared the automatically segmented images with manual measurements of fungal colony size in each picture. Once the images were loaded into the software *ImageJ*, the manual measurements took on average 4 minutes and 52 seconds per isolate and growth condition (with 22 images measured on average per isolate) (Table 1). The growth data obtained by manual measurements was similar to the automatic segmentation (Fig. 3), and there were no significant differences in slope estimates based on the data obtained with the two methods (Table 2). The pairwise difference between manual and automatic annotation was on average 12.8% (Standard Error of the mean = 3.7%) (Table 2), which is not larger than the variation inherent between consecutive manual measurements of the same image.

**Table 1.** Time used on measuring manual segmentation using ImageJ.

| Isolate | pH | Time used on<br>manual measurement | Number of<br>images measured |
| --- | --- | --- | --- |
| C005 | 3 | 5 min. and 35 sec. | 36 |
| C006 | 3 | 5 min. and 38 sec. | 38 |
| C008 | 3 | 5 min. and 3 sec. | 38 |
| C009 | 3 | 4 min. and 49 sec. | 38 |
| C005 | 5.6 | 3 min. and 47 sec. | 8 |
| C006 | 5.6 | 3 min. and 43 sec. | 14 |
| C008 | 5.6 | 4 min. and 57 sec. | 21 |
| C009 | 5.6 | 5 min. and 25 sec. | 21 |
|  |  | Mean: 4 min. and 52 sec. | Mean: 22 images |

**Table 2:** Overview of datapoints used to estimate and statistically compare the steepest slope obtained from manually and automatically segmented data respectively. Timepoints refer to the placement of a minimum of four consecutive datapoints yielding the steepest slope.

| Isolate | pH | Timepoints<br>(Hours) | Slope manual<br>(pixels/hour) | Slope auto<br>(pixels/hour) | Slope<br>difference (%) | Pairwise slope<br>comparison |
| --- | --- | --- | --- | --- | --- | --- |
| C005 | 3 | 22-40 | 337 | 343 | 1.76 | $t_{16} = 0.16$<br>$p_{\text{BH-adj.}} = 0.9915$ |
| C006 | 3 | 26-48 | 595 | 746 | 22.52 | $t_{20} = 2.55$<br>$p_{\text{BH-adj.}} = 0.1512$ |
| C008 | 3 | 20-28 | 246 | 226 | 8.47 | $t_6 = 0.61$<br>$p_{\text{BH-adj.}} = 0.8990$ |
| C009 | 3 | 20-28 | 363 | 301 | 18.67 | $t_5 = 2.10$<br>$p_{\text{BH-adj.}} = 0.3580$ |
| C005 | 5.6 | 20-26 | 3350 | 2457 | 30.76 | $t_4 = 1.86$<br>$p_{\text{BH-adj.}} = 0.3621$ |
| C006 | 5.6 | 20-26 | 2168 | 2173 | 0.23 | $t_4 = 0.01$<br>$p_{\text{BH-adj.}} = 0.9915$ |
| C008 | 5.6 | 20-30 | 1469 | 1364 | 7.41 | $t_8 = 0.42$<br>$p_{\text{BH-adj.}} = 0.9145$ |
| C009 | 5.6 | 20-30 | 1571 | 1385 | 12.58 | $t_8 = 0.88$<br>$p_{\text{BH-adj.}} = 0.8086$ |

**Figure 3.**
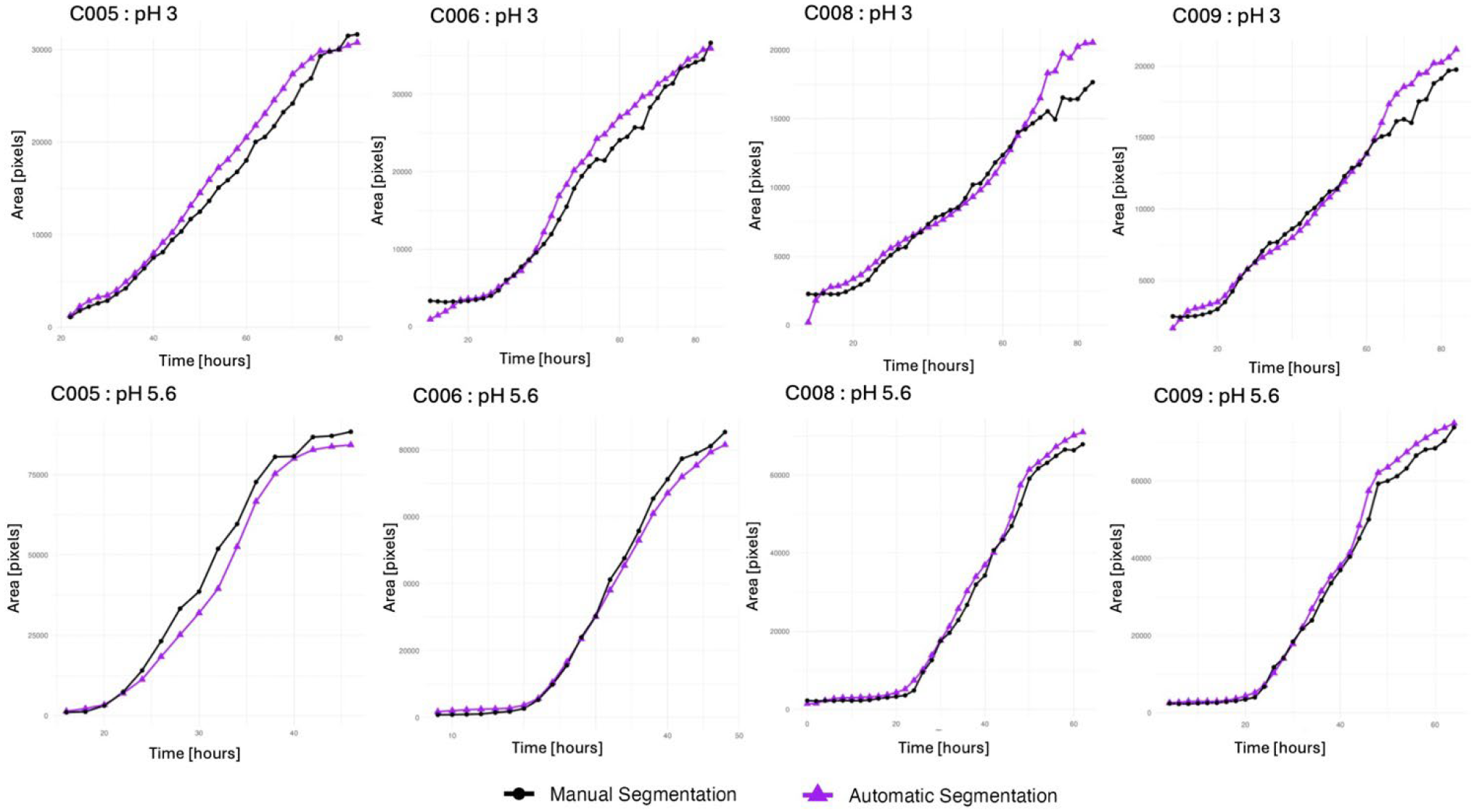
Comparison of area over time for manual measurements (black circle) and automatic segmentations (purple triangle) carried out on images in Test set 1 (one replicate of each combination of isolate and pH level). Statistical analysis of slope differences are provided in Table 2.

In general, isolates grown on standard PDA with a pH level of 5.6 grew faster than isolates grown on PDA with a pH level adjusted to pH 3 (Fig. 4). Moreover, the two *C. gloeosporioides s. lat*. isolates grew faster than the *C. acutatum s. lat*. isolates, and the variability between the group means is significantly greater than the variability within the groups (one-way ANOVA: F_7,40_ = 72.98, *p <* 2e-16). Using TukeyHSD post-hoc tests we further observe that the eight isolate-pH combinations fall into three groups, that are significantly different from each other (Fig. 4, See supplementary material). The maximum growth rates of all four isolates, when grown at pH 3 (group A), are not significantly different from each other, while they are significantly different from all isolates grown on standard PDA with pH 5.6. When grown on standard PDA, the isolates fall into groups according to species complexes (Fig. 4): The maximum growth rates of the two *C. gloeosporioides s. lat*. are not significantly different from each other (group B), and the maximum growth rates of the two *C. acutatum s. lat*. are not significantly different from each other (group C). Simultaneously, isolates of the two species complexes are significantly different from each other and from all isolates grown at pH 3. The intra-isolate (replicate) variance is higher at pH 5.6 compared to pH 3, especially for the two *C. gloeosporioides s. lat*. isolates C005 and C006 (Fig. 4, See supplementary material).

**Figure 4.**
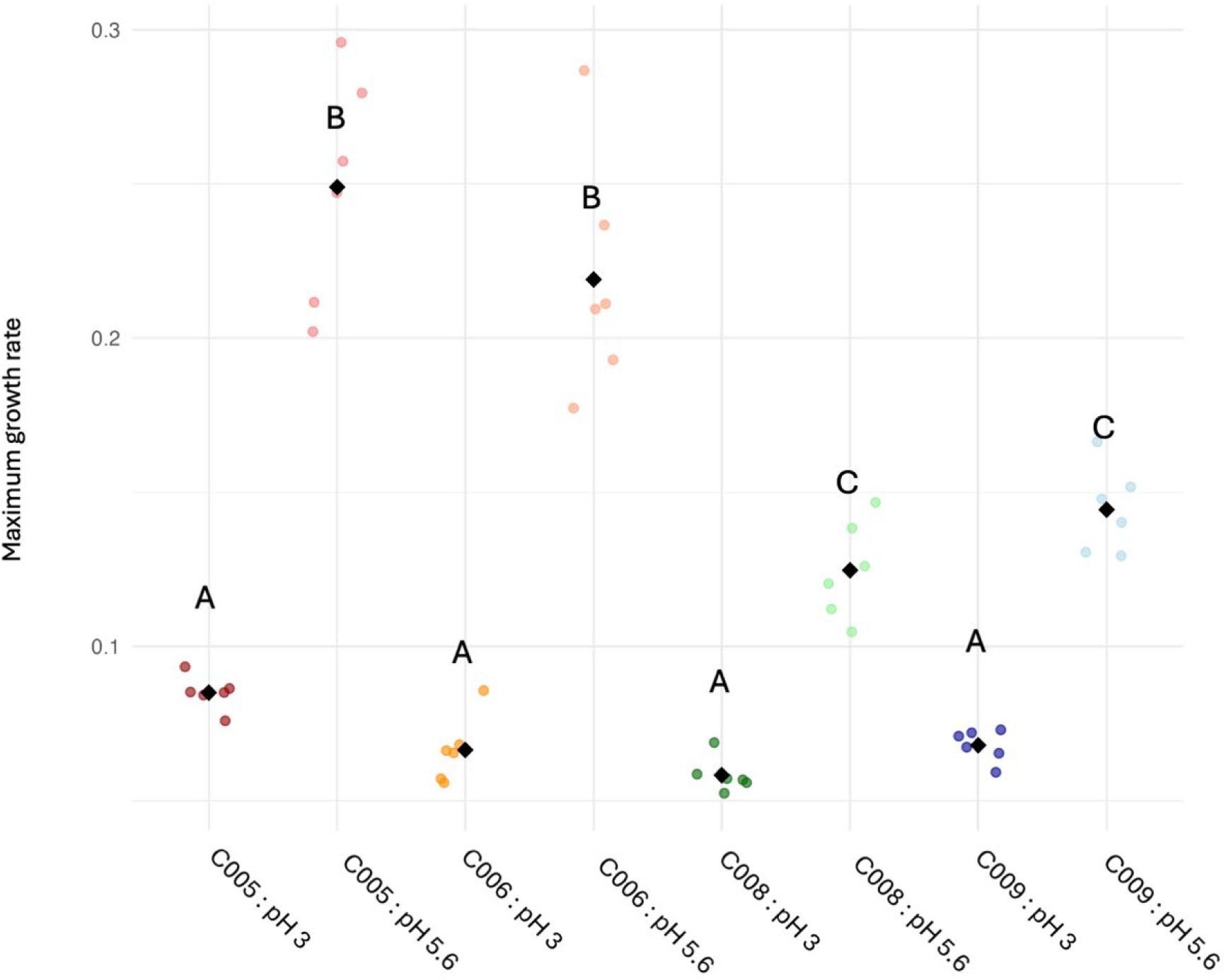
Maximum growth rate determined from automatically segmented images for each combination of isolate and pH level. The mean for each combination is shown as black squares. A total of 2,016 images (four fungal isolates at two pH levels in six technical replicates) were analyzed. Capital letters A, B, and C show significant groupings based on TukeyHSD post-hoc tests.

## Discussion

Here we developed a methodological analysis pipeline to automatically segment time-lapse images of growing fungal cultures on solid media. With limited hands-on time required by the experimenter to train, verify and improve the deep-learning model to automatically segment the images. Under our conditions the method is equally as good as traditional manual image measurements, and the automatic segmentation based on the deep learning model provides data as accurate as traditional manual methods. Our method is thus able to clearly distinguish growth rate of our isolate-pH combinations from each other. Some variation between replicate wells and images of growing fungi is inherent, and this variation would most likely also have been observed for manual measurements. This variation is often due to high transparency of the mycelium especially at the edge of growing colonies that make measurements challenging. A benefit of the automatic segmentation is that the same bias is applied to all images, while this might vary in manual segmentation. The main advantage of our method is speed. It took less than five minutes to segment the 2,016 images automatically once the model was trained. Based on time measurements done during manual segmentation, this would have taken approximately 4 hours to manually segment. Time spent on manual measurements increases linearly with the number of isolates, replicates and/or treatments (e.g., pH level) and thus quickly becomes a bottleneck or limitation in experimental set-ups. This problem is circumvented with automatic segmentation as the analysis can run without the need of human interaction and computational power can be increased to reduce run-time.

We assessed growth of four isolates of *Colletotrichum* fungi as an example of a filamentous ascomycete fungus. Our obtained results are in correspondence with existing knowledge of *C. gloeosporioides s. lat*. generally having a higher maximum growth rate than *C. acutatum s. lat*. and the influence of pH on fungal growth (Greer et al. 2011, Mcgovern et al. 2012, Smith & Black 1990). However, it is important to stress that the iterative nature of how the automatic segmentation model is trained implies that the method presented here is equally applicable to other fungal species or fungi with markedly different morphologies. Because the image segmentation is consistently verified by the experimenter (Smith et al. 2022), the limiting factor is likely the contrast in the images between the fungus and the background. For fungi with very sparse growth and hyphae with the same color as the media, applying automatic image segmentation is likely going to be more challenging like if manual measurements were used.

Here we used an automated image acquisition system to obtain images, but any standardized method to obtain images will work. While limited variation in lighting and composition (such as centralizing the fungus culture in the images) is beneficial for training the model, it is not a necessity. The model can be effectively trained and correctly segment image pixels as either fungal mycelium or background, regardless. This image-based method allows for the capturing of phenotypic traits of fungi growing on solid media that would not be detected by OD-based measurements of fungi growing in liquid media (Slowik et al. 2023). It would for example be possible to analyze the shape pattern of the growing mycelium by calculating *compactness* defined as area divided by length of the colony edge, or variation in *texture* by analyzing variation in entropy of the fungus-segmented area of images. Liquid cultivation and OD-based methods using microtiter plates offer high-throughput and the 24-well plates used here begin to approach the same level of throughput. In addition, the image-based method provides a means to assess contamination or other abnormalities that can be visually inspected on the images of outlier datapoints, which would not be possible with OD-based measurements of microtiter plates. Finally, measuring growth on solid media versus submerged liquid growth also often allow onset of sporulation to be captured, if spores can be distinguished from the mycelium, for example based on color.

## Conclusion

Our results demonstrate that the semi-automatic *RootPainter* segmentation method resulted in growth curves that were indistinguishable from those resulting from manual area estimations. After less than two hours of interactive training, *RootPainter* automatically segments the full image collection. This saves hours of manual measurements and thereby allows much higher time resolution for more accurate profiling of growth curves. The ease and speed whereby a specific deep learning model can be trained and used on new data ensures that the method presented here can be adjusted for any fungal species where filamentous growth on solid media can be captured by digital images. As such this method is widely applicable across fungal species and requires little specialized programming skills by the experimenter making it more accessible to experimental mycologists.

## Acknowledgments

The authors thank Italo Delalibera Jr. and Natasha Iwanicki for providing the initial fungal isolates used to develop and test this method. HHDFL and this research were supported by a Carlsberg Foundation Semper Ardens: Accelerate grant (CF20-0609).

## Author contribution

TK, EBD and HDFL conceived the idea. TK and HDFL designed the experimental setup. TK and EBD designed the deep learning model methodology. TK, EBD and HDFL designed the analysis pipeline. TK planned and carried out the experimental work, model training and implementation of the analysis pipeline. TK led the writing and wrote the first draft of the manuscript with input from HDFL and EBD. TK, EBD and HDFL wrote and edited later manuscript drafts and all authors gave final approval for publication.

