## Supplementary file for "A high-throughput method for measuring fungal growth rate on solid media using automated imaging and deep learning"

### Setting up

Loading necessary packages and setting working directory.

```
# load packages
library(growthrates)
library(ggplot2)
library(dplyr)
library(tidyr)
library(stringr)
library(gridExtra)
library(dpseg)
library(lsmeans)
```

Setting working directory.

```
setwd("~/Desktop/deepLearningMethodPaper/fungalGrowthRate_deepLearning_forPublication/")
```

### Comparing manual and automatic methods

#### Manual and automatic segmentation of Dataset1

I have manually measured the diameter twice c005, c006, c008 and c009 for both pH 3 and pH 5.6 in ImageJ. Note that it was difficult to manually measure at pH 5.6 due to high degree of transparency of mycelium.

I have collected all the measurements from ImageJ in one .csv file, that I load in as a dataframe. I further calculate the average diameter for each image and calculate the area from this. I make sure the dataframe only contains the needed columns. And in the end I multiple the time with 2 in order to get the time unit in hours (as pictures were taken every second hour).

```
# loading in the manual measurements
df_manual <- read.csv("./manSegmImageJ_c5c6c8c9_ph3_pHdot6.csv")
df_manual <- df_manual %>%
  select(Label, Length)

#split label
df_manual <- df_manual %>%
  mutate(
    Isolate = str_extract(Label, "c\\d{3}"),
    Time = str_extract(Label, "T\\d+") %>% str_replace("T", ""),
    Time = as.integer(Time),
    pH = str_extract(Label, "(?i)ph([0-9]+(?:dot[0-9]+)?)")) %>%
    mutate(pH = sub("(?i)ph", "", pH)) %>%
    mutate(Well = str_extract(Label, "_[A-Z][0-9]_")) %>%
    mutate(Well = sub("_", "", Well)) %>%
    mutate(Well = sub("_", "", Well)
  )

# make average length
# Group by "pH", "Isolate", and "Time", and calculate the average of "Length"
df_manual <- df_manual %>%
  group_by(pH, Isolate, Time, Well) %>%
```

```r
  summarise(Length = mean(Length), .groups = 'drop')

#calculate area from diameter (length)
# Calculating an area based on the diameter (column Length), under the assumption of circular growth:
# area = r^2 * pi
df_manual <- df_manual %>%
  mutate(Area = pi * (Length / 2)^2)

# make sure its only same columns as in auto
df_manual <- df_manual %>%
  select(pH, Well, Isolate, Time, Area)

#multiplying all T timepoints with two as it is every second hour
df_manual <- df_manual %>%
  mutate(Time = Time * 2)
```

Then I load in the automatic measurements as a dataframe. When using the model for segmentation, sometimes it finds more than one segment. I make sure that there is only one area per time to work around this issue.
Here I also multiple the time with 2 in order to get the time unit in hours (as pictures were taken every second hour).

```r
# loading in the automatic measurements
df_auto <- read.csv("./segmc5c6c8c9_ph3-ph5dot6.csv")

df_auto <- df_auto %>%
  select(file_name, area)

#split file_name
df_auto <- df_auto %>%
  separate(file_name, into = c("pH", "Well", "Isolate", "Time"), sep = "_")

#change so pH and T is just a number
df_auto <- df_auto %>%
  mutate(
    Time = sub("T", "", Time),
    Time = as.integer(Time),
    pH = sub("(?i)ph", "", pH)
    )

# Only keep highest area as the model sometimes makes more than one segment
# Group by Isolate, Well, pH, and Time, and filter to keep the row with the highest area
df_auto <- df_auto %>%
  group_by(Isolate, Well, pH, Time) %>%
  filter(area == max(area)) %>%
  ungroup()

#multiplying all T timepoints with two as it is every second hour
df_auto <- df_auto %>%
  mutate(Time = Time * 2)
```

Then I combine the automatic and manual measurements in one dataframe, I ensure time is an integer and then I plot area over time for each combination of isolate:pH.

```r
# Check column names
colnames(df_auto)
```

```
## [1] "pH"      "Well"    "Isolate" "Time"    "area"
```

```r
colnames(df_manual)
```

```
## [1] "pH"      "Well"    "Isolate" "Time"    "Area"
```

```r
# Combined dataframe for manual and automatic segmentation
df_segManAut_combined <- merge(df_auto, df_manual, by = c("Time", "Well", "Isolate", "pH"))

#Make time integer
df_segManAut_combined <- df_segManAut_combined  %>%
  mutate(Time = as.integer(Time))

# Get unique combinations of isolate and ph and order by isolate
combinations <- df_segManAut_combined %>%
  select(Isolate, pH) %>%
  distinct() %>%
  arrange(Isolate, pH)

# Create a list to store the plots
plot_list <- list()

# Plotting for each combination of isolate and ph
for(i in 1:nrow(combinations)) {
  isolate <- combinations$Isolate[i]
  ph <- combinations$pH[i]

  df_subset <- df_segManAut_combined %>% filter(isolate == Isolate, ph == pH)

  p <- ggplot(df_subset, aes(x = Time)) +
    geom_line(aes(y = area, color = "Automatic Segmentation"), size = 1) +
    geom_line(aes(y = Area, color = "Manual Segmentation"), size = 1) +
    geom_point(aes(y = area, color = "Automatic Segmentation"), shape = 17, size = 3) +  # Change shape
    geom_point(aes(y = Area, color = "Manual Segmentation"), size = 2) +
    labs(title = paste("Isolate:", isolate, "pH:", ph),
         x = "Time",
         y = "Area") +
    scale_color_manual(values = c("Automatic Segmentation" = "purple", "Manual Segmentation" = "black"))
    theme_minimal()

   # Save the individual plot as a PNG file
  filename <- paste0("plot_compare_manAut_pH_", ph, "_Isolate_", isolate, ".png")
  ggsave(filename, plot = p, width = 8, height = 6)

    # Add the plot to the list
  plot_list[[i]] <- p

  print(p)
}
```

```
## Warning: Using 'size' aesthetic for lines was deprecated in ggplot2 3.4.0.
## i Please use 'linewidth' instead.
## This warning is displayed once every 8 hours.
## Call 'lifecycle::last_lifecycle_warnings()' to see where this warning was
## generated.
```

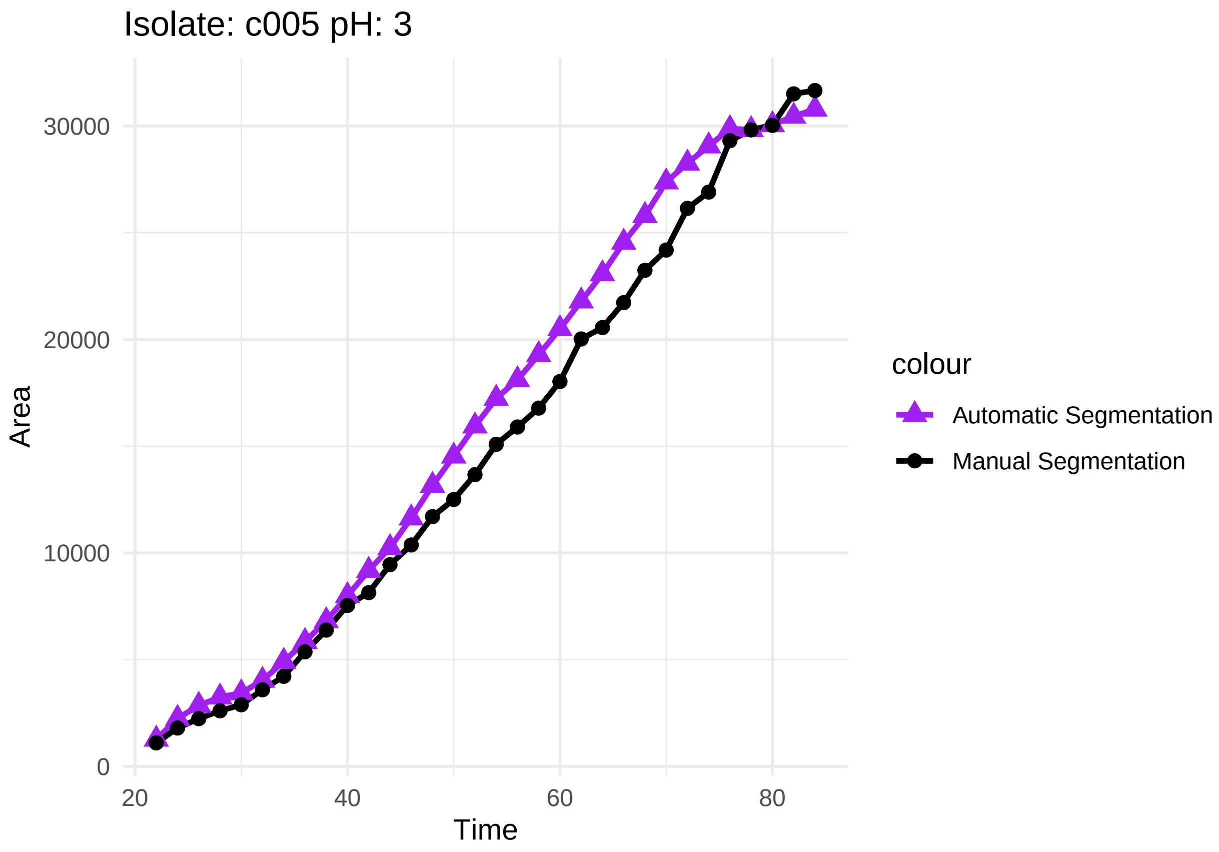

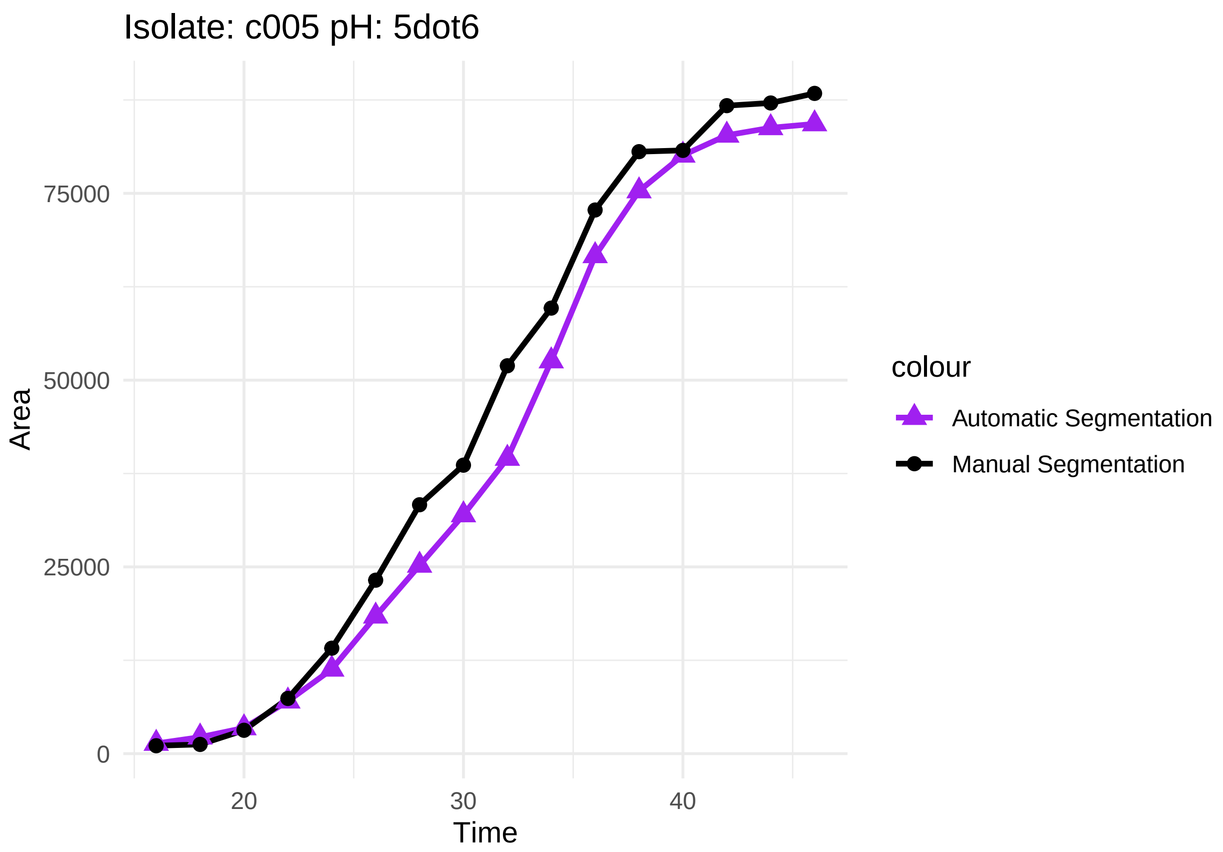
Isolate: c005 pH: 5dot6
Area
75000
50000
25000
0
20
30
40
Time
colour
Automatic Segmentation
Manual Segmentation

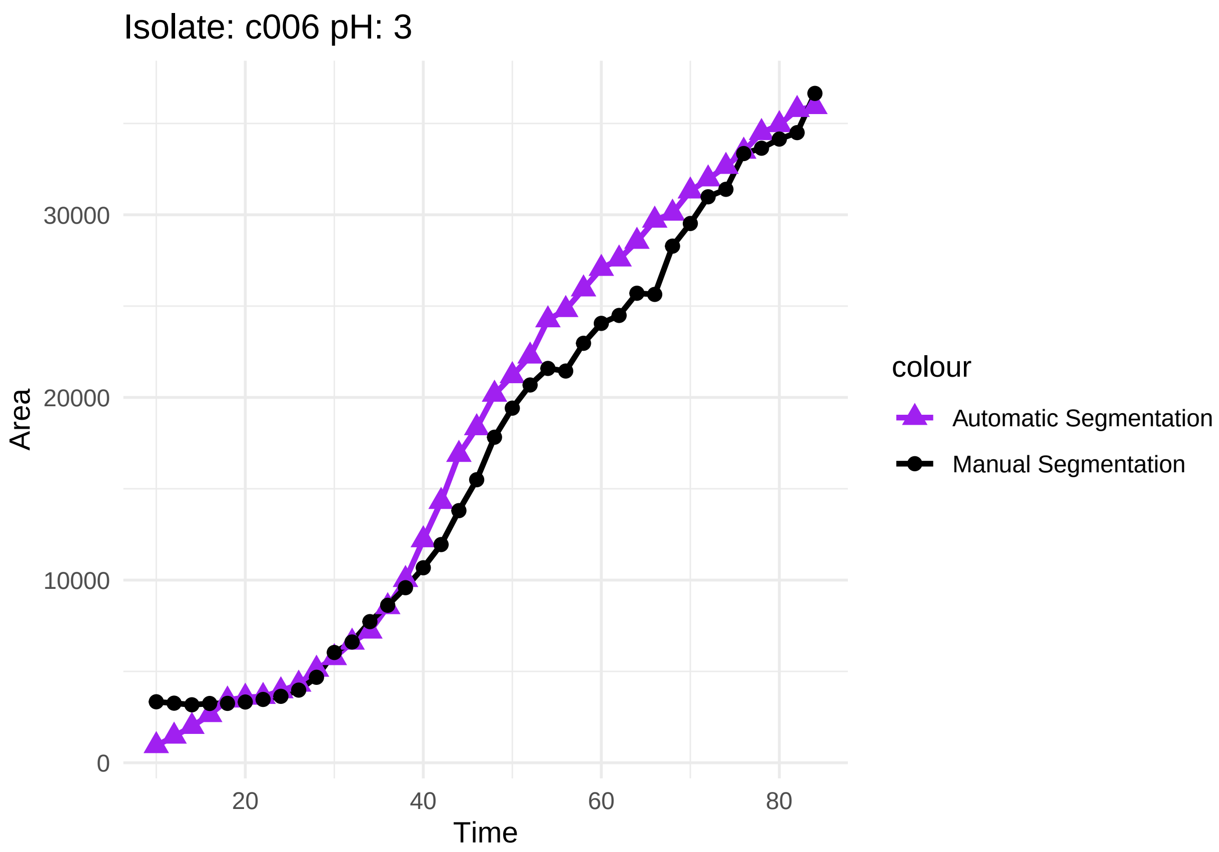
Isolate: c006 pH: 3
Area
30000
20000
10000
0
20
40
60
80
Time
colour
Automatic Segmentation
Manual Segmentation

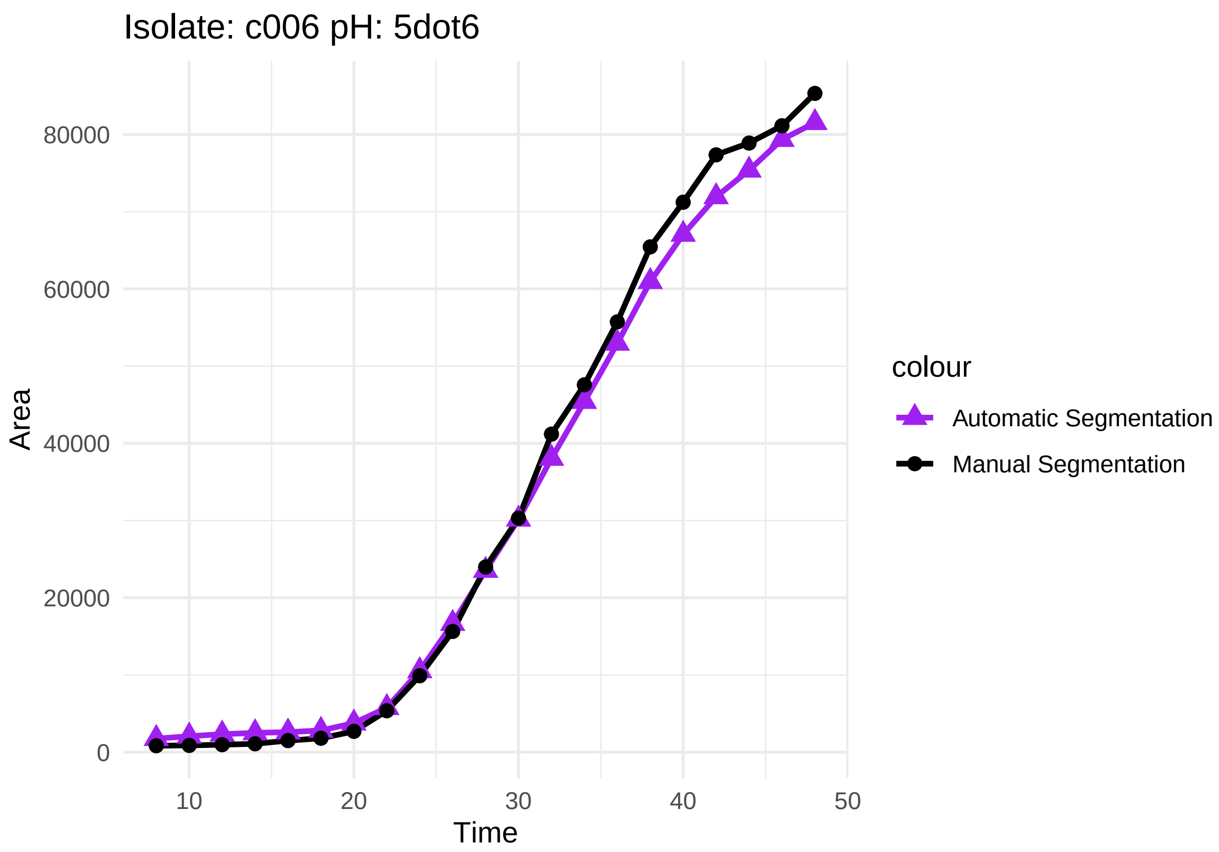
Isolate: c006 pH: 5dot6
Area
80000
60000
40000
20000
0
Time
10
20
30
40
50
colour
Automatic Segmentation
Manual Segmentation

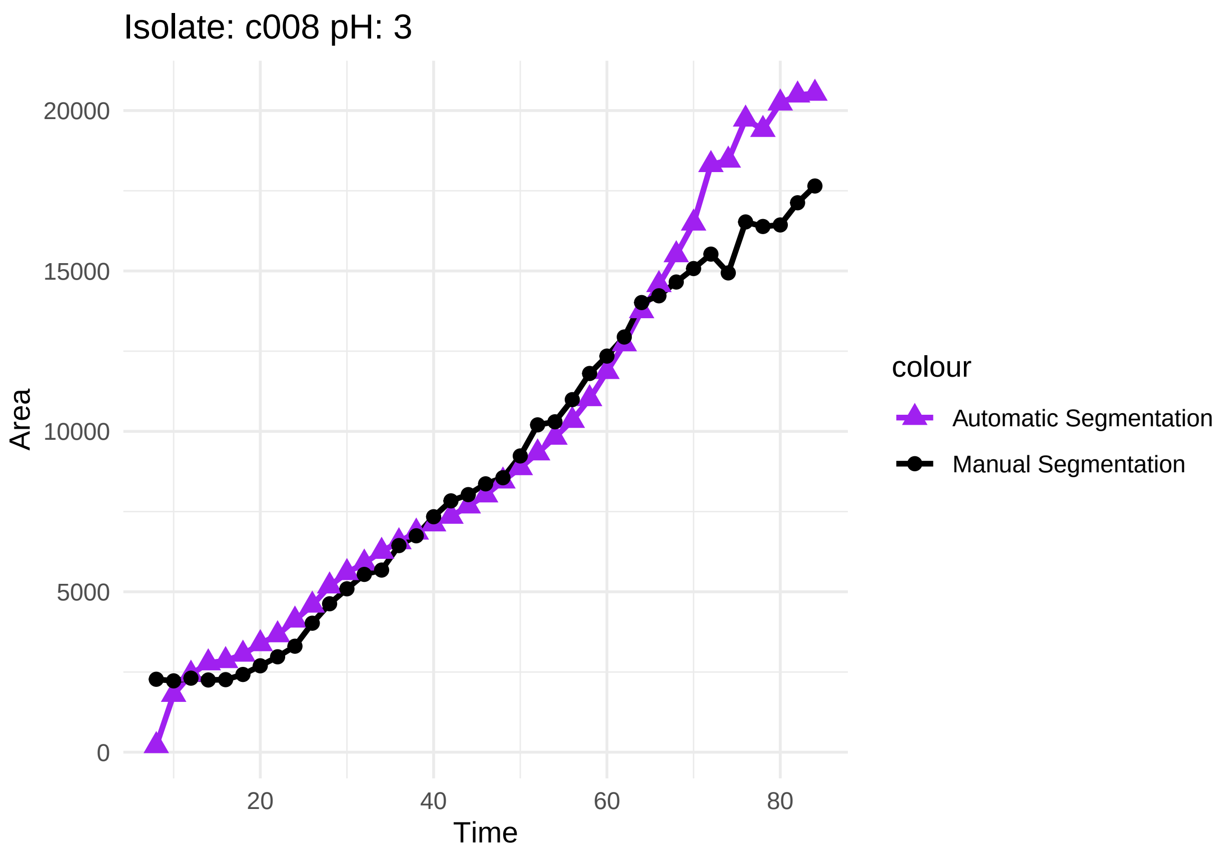
Isolate: c008 pH: 3
20000
15000
10000
5000
0
Area
20
40
60
80
Time
colour
Automatic Segmentation
Manual Segmentation

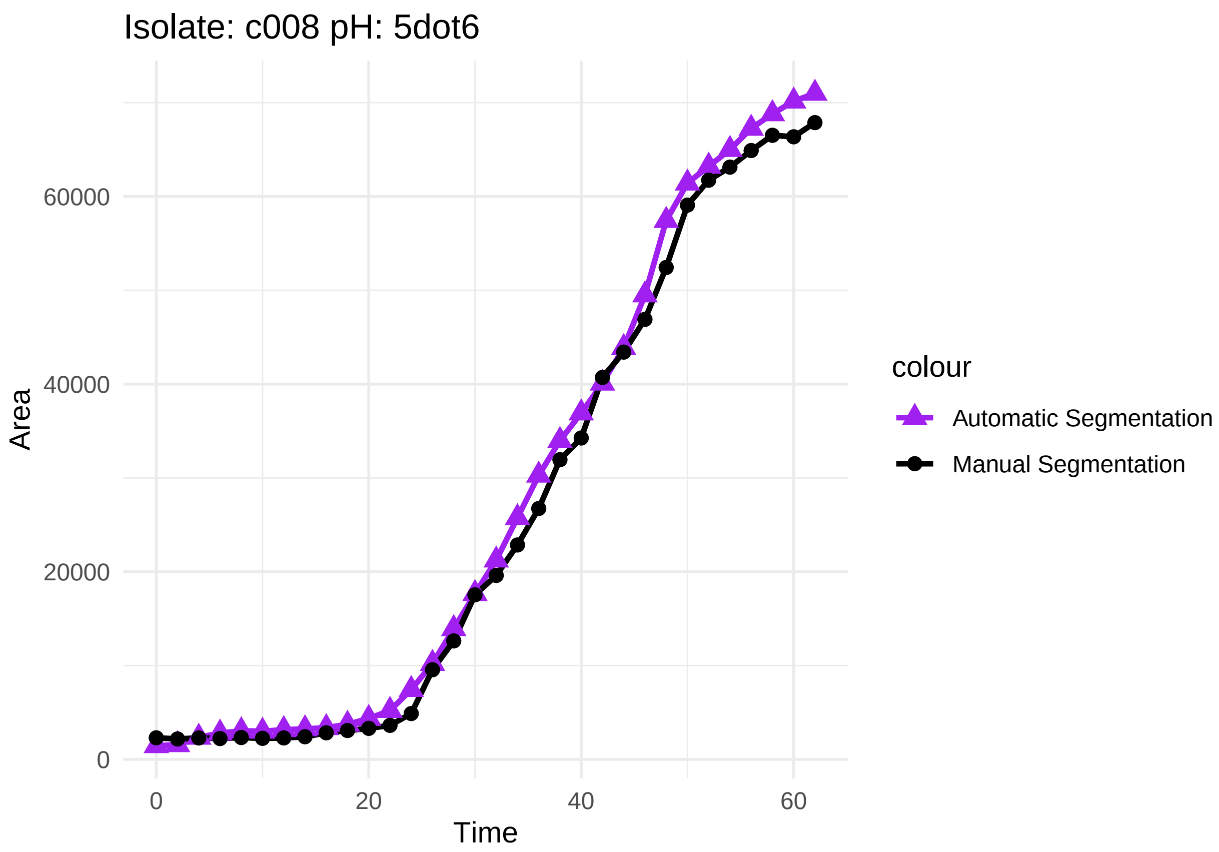

Isolate: c008 pH: 5dot6
Area
60000
40000
20000
0
0
20
40
60
Time
colour
Automatic Segmentation
Manual Segmentation

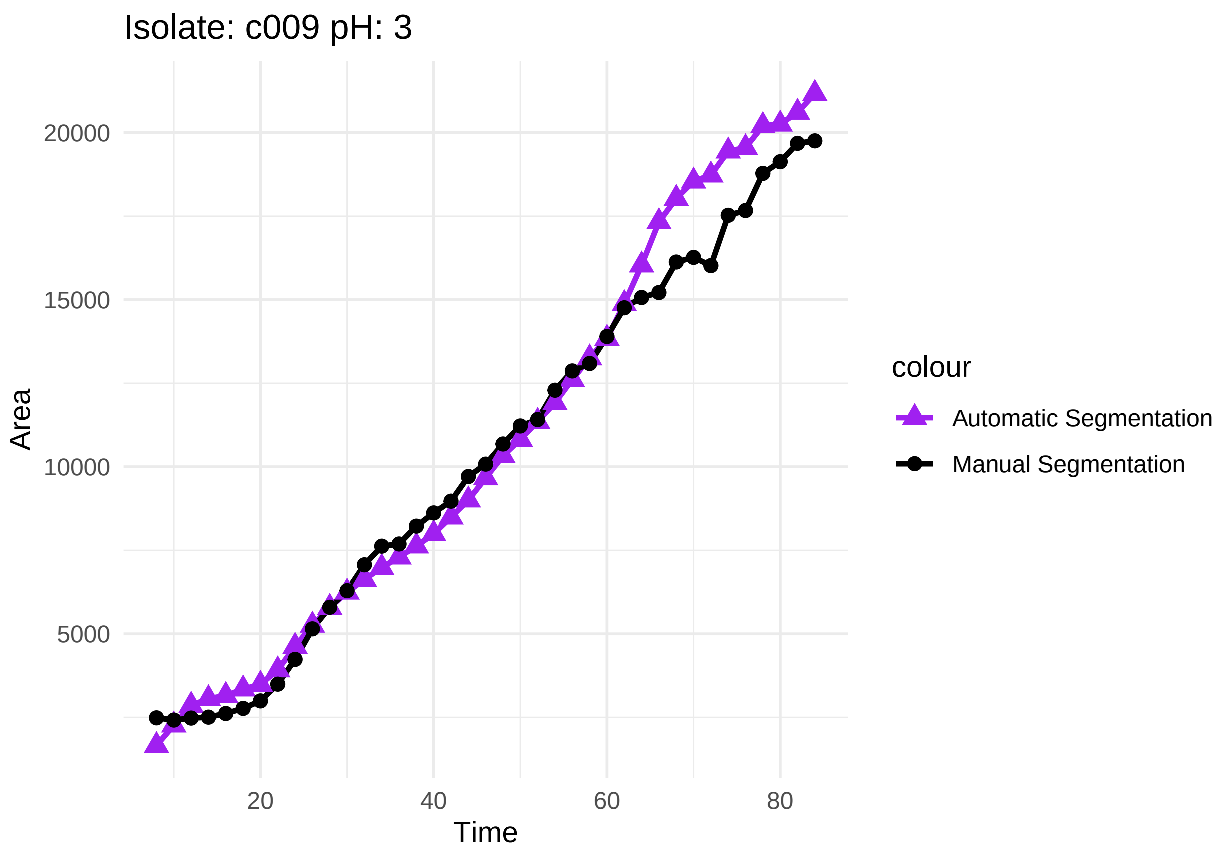
Isolate: c009 pH: 3
20000
15000
10000
5000
Area
20
40
60
80
Time
colour
Automatic Segmentation
Manual Segmentation

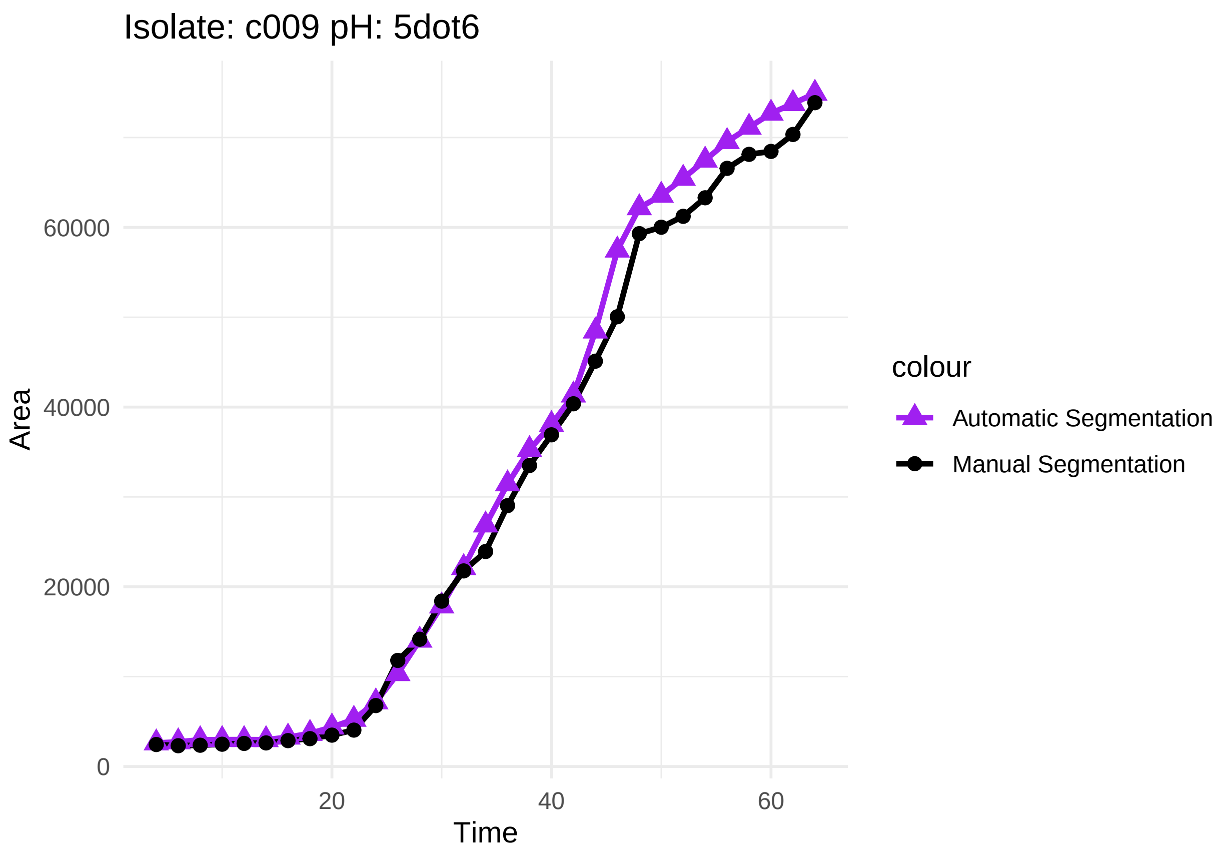

Here the plots as above, but with natural log transformed area:

```
# Create a list to store the plots
plot_list <- list()

# Plotting for each combination of isolate and ph
for(i in 1:nrow(combinations)) {
  isolate <- combinations$Isolate[i]
  ph <- combinations$pH[i]

  df_subset <- df_segManAut_combined %>% filter(isolate == Isolate, ph == pH)

  p <- ggplot(df_subset, aes(x = Time)) +
    geom_line(aes(y = area, color = "Automatic Segmentation"), size = 1) +
    geom_line(aes(y = Area, color = "Manual Segmentation"), size = 1) +
    geom_point(aes(y = area, color = "Automatic Segmentation"), shape = 17, size = 3) +  # Change shape
    geom_point(aes(y = Area, color = "Manual Segmentation"), size = 2) +
    labs(title = paste("Isolate:", isolate, "pH:", ph),
         x = "Time",
         y = "Area") +
    scale_color_manual(values = c("Automatic Segmentation" = "purple", "Manual Segmentation" = "black"))
    theme_minimal()

   # Save the individual plot as a PNG file
  #filename <- paste0("plot_pH_", ph, "_Isolate_", isolate, ".png")
  #ggsave(filename, plot = p, width = 8, height = 6)
```

```
    # Add the plot to the list
  plot_list[[i]] <- p

  print(p)
}
```

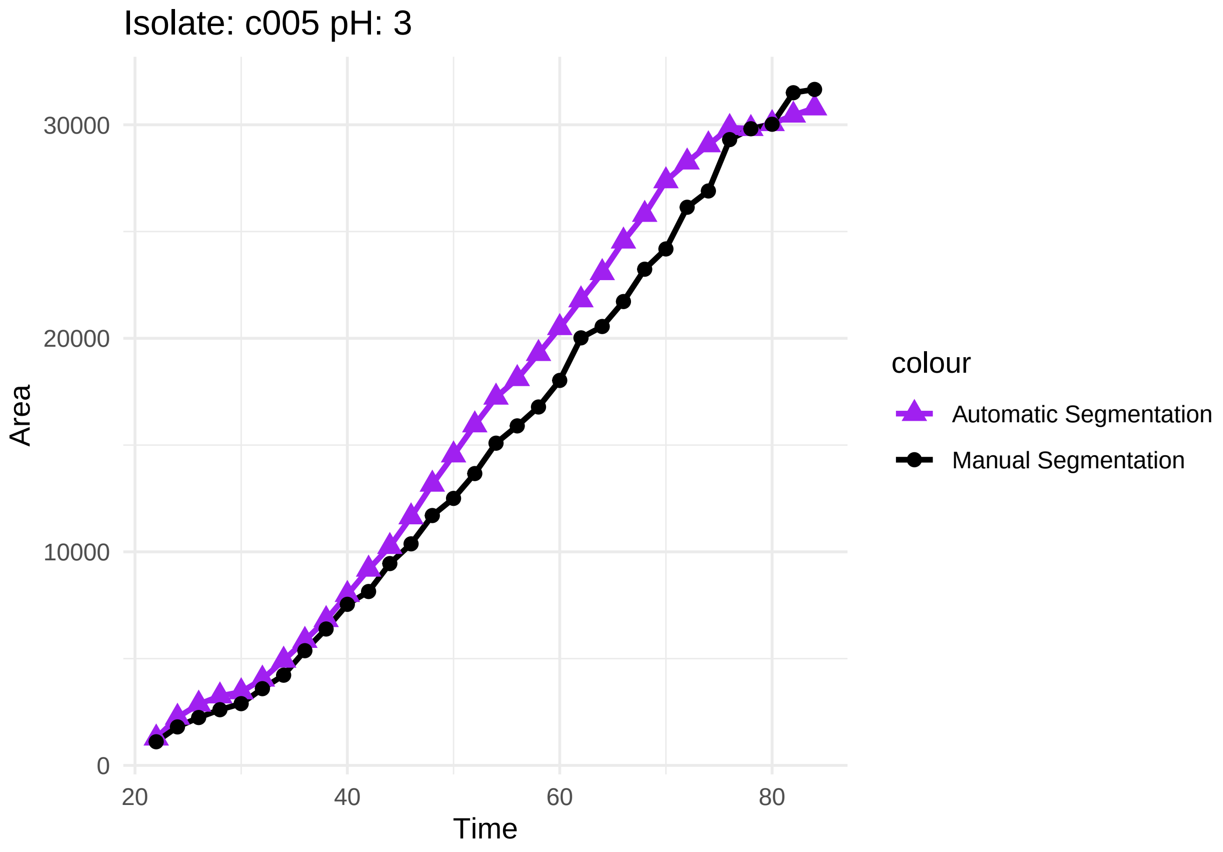

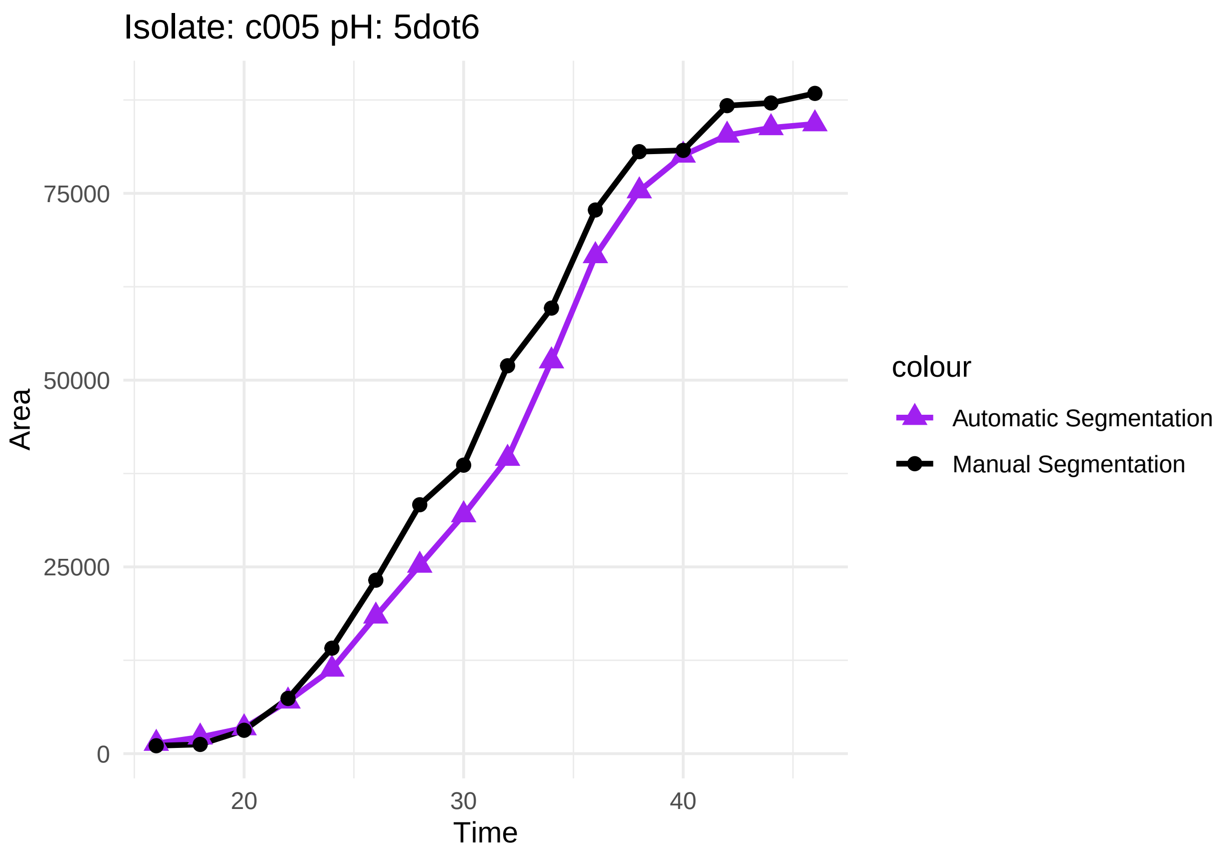

Isolate: c005 pH: 5dot6
Area
75000
50000
25000
0
20
30
40
Time
colour
Automatic Segmentation
Manual Segmentation

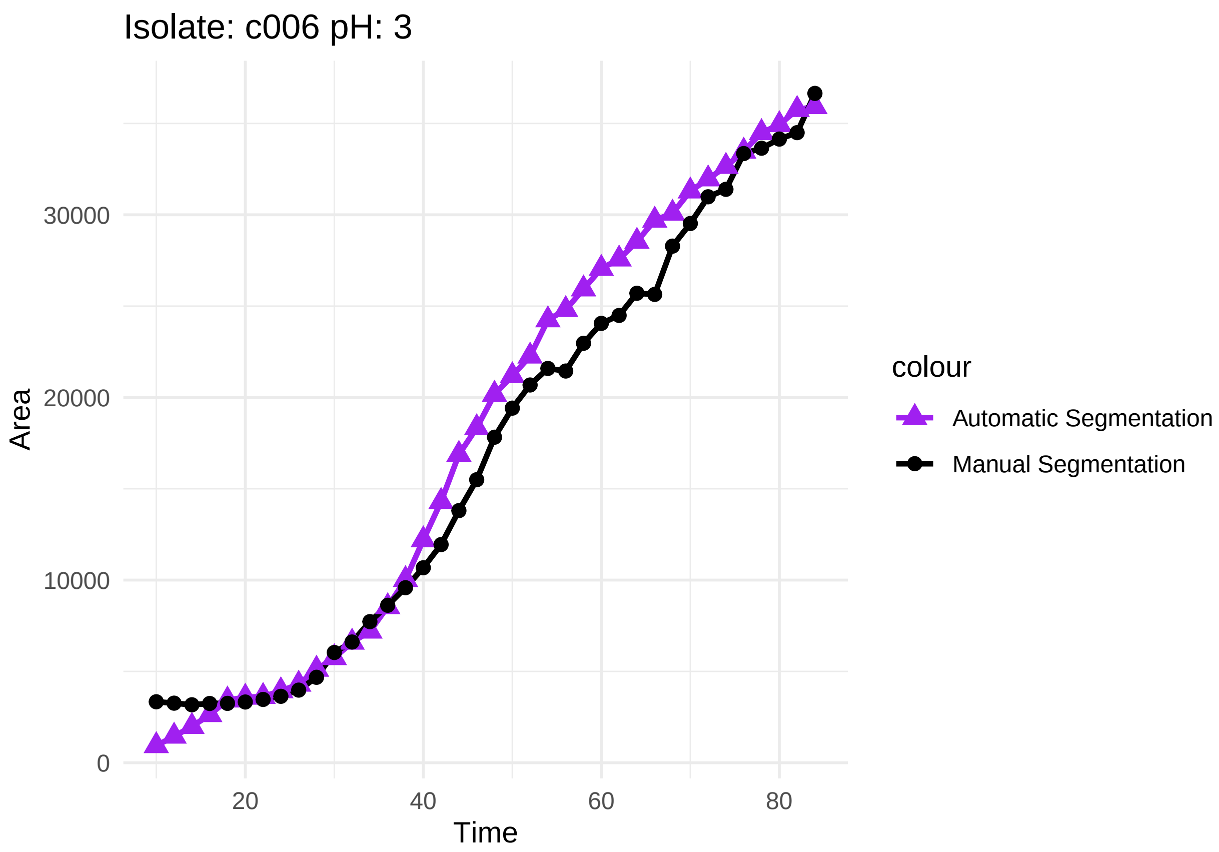
Isolate: c006 pH: 3
Area
Time
0
10000
20000
30000
20
40
60
80
colour
Automatic Segmentation
Manual Segmentation

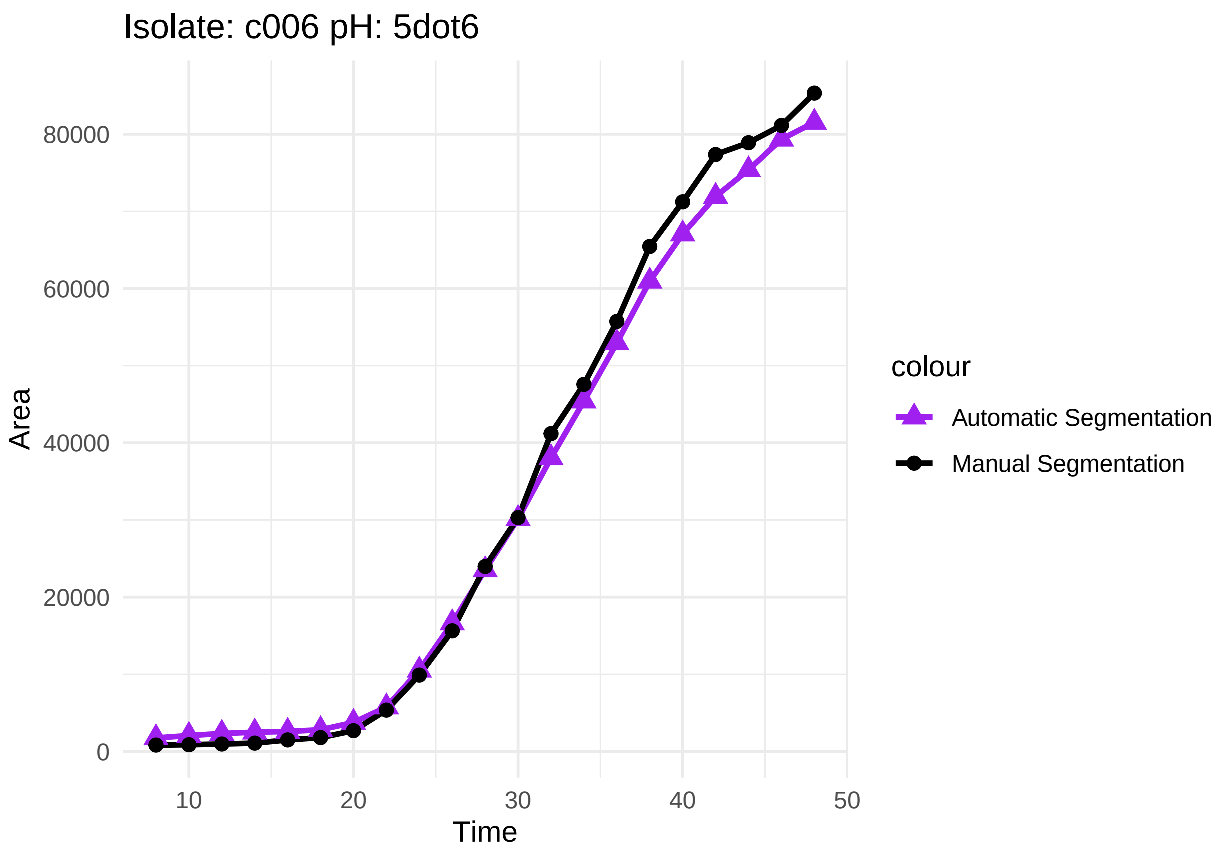
Isolate: c006 pH: 5dot6
80000
60000
40000
20000
0
Area
10
20
30
40
50
Time
colour
Automatic Segmentation
Manual Segmentation

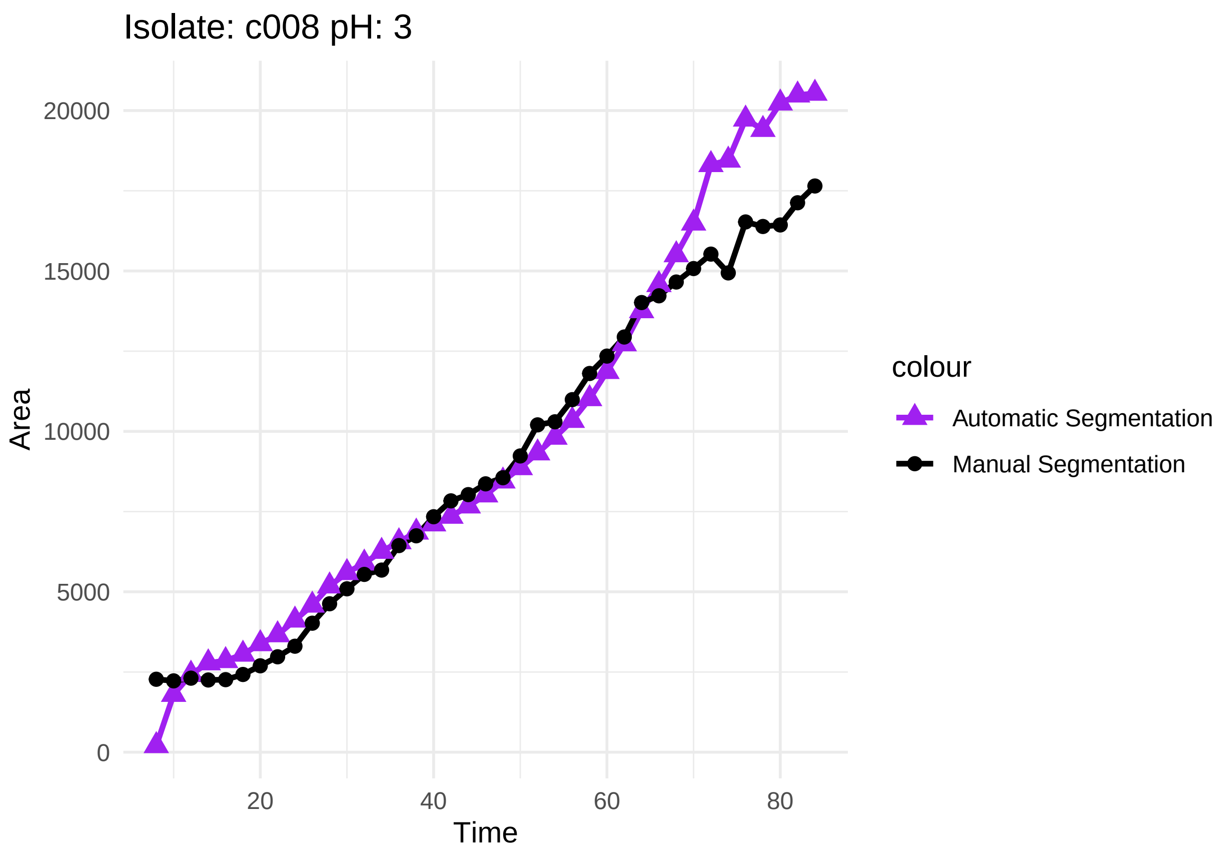

Isolate: c008 pH: 3
20000
15000
10000
5000
0
Area
20
40
60
80
Time
colour
Automatic Segmentation
Manual Segmentation

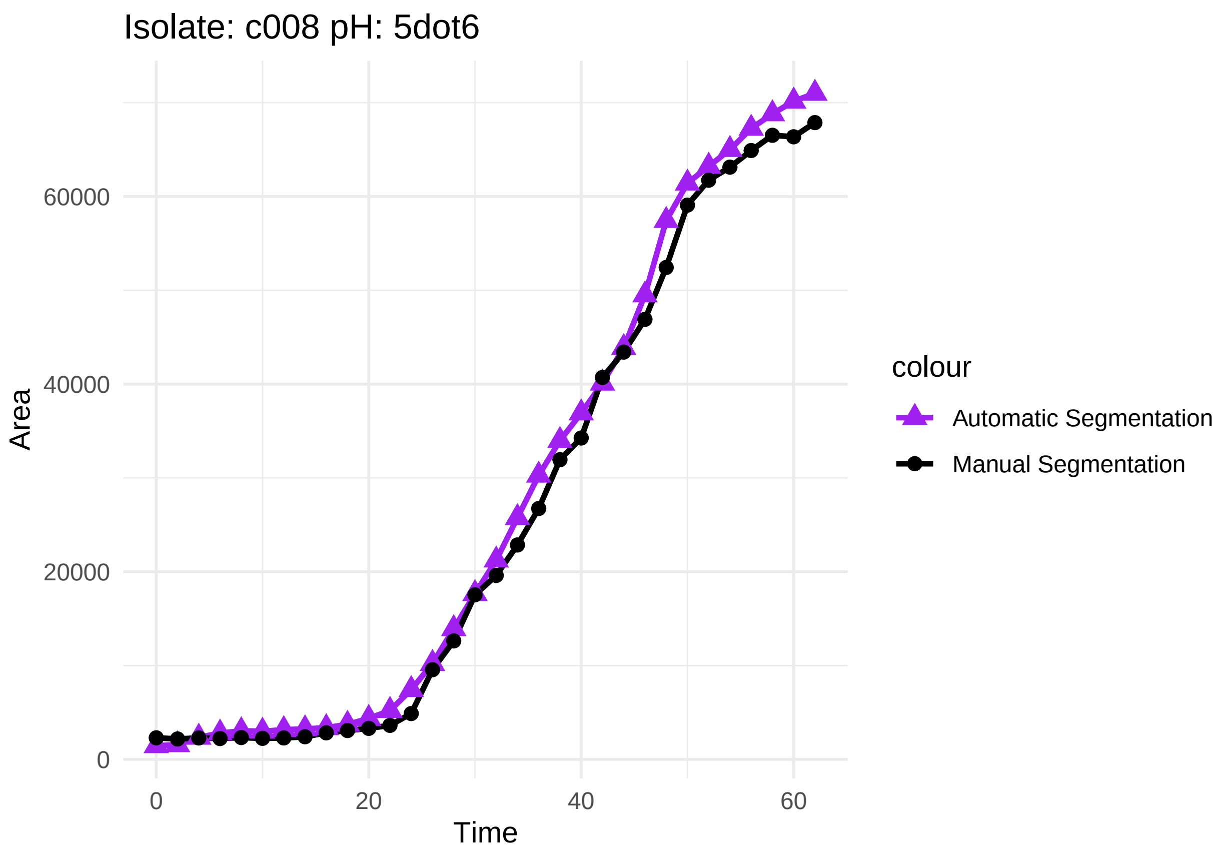
Isolate: c008 pH: 5dot6
Area
60000
40000
20000
0
0
20
40
60
Time
colour
Automatic Segmentation
Manual Segmentation

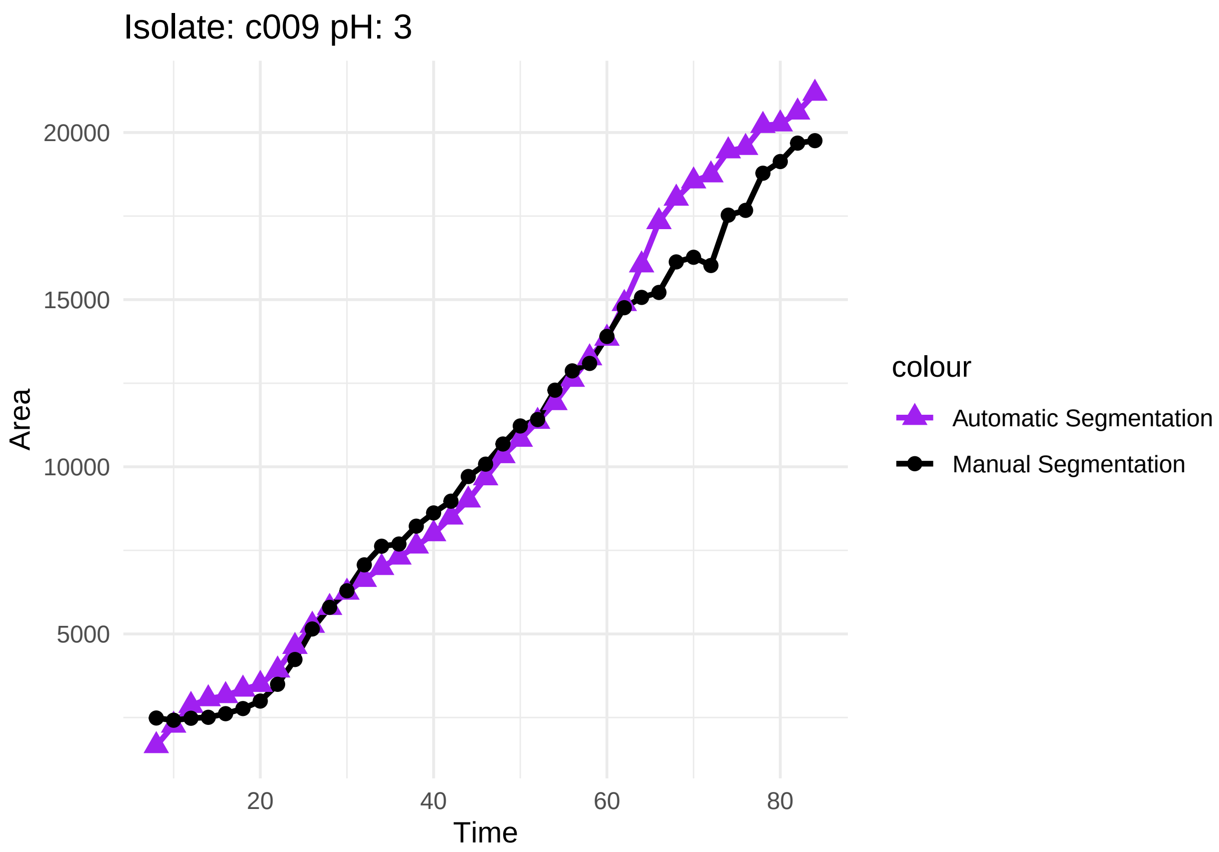
Isolate: c009 pH: 3
Area
20000
15000
10000
5000
20
40
60
80
Time
colour
Automatic Segmentation
Manual Segmentation

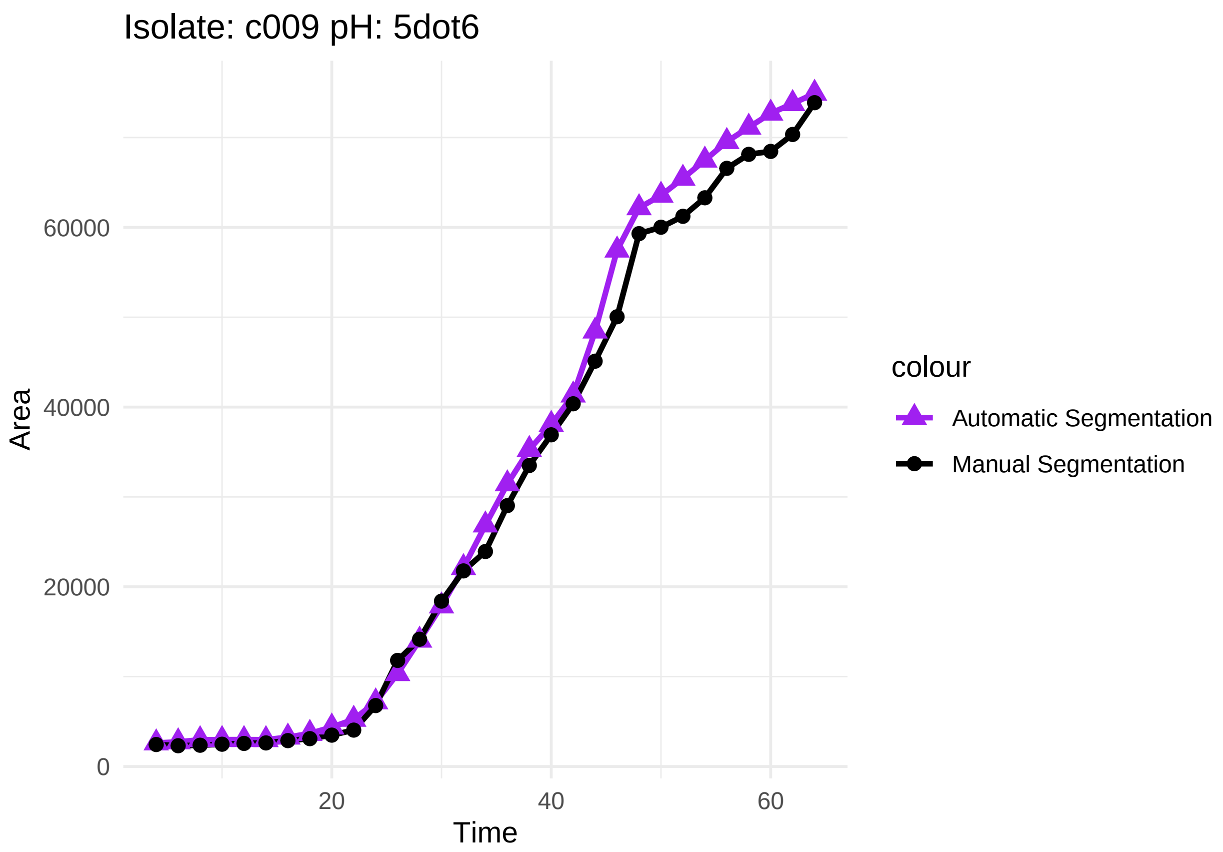

I first tweak the dataframe that contains both manual and auto data, so column Area contains for both methods, and column Isolate tells if it is auto or manual. And I make a new column logArea where I have natural log transformed area.

```
# Create a new dataframe for the 'area' entries with modified 'Isolate' column
df_area_auto <- df_segManAut_combined %>%
  select(Time, Well, Isolate, pH, area) %>%
  rename(Area = area) %>%
  mutate(Isolate = paste0(Isolate, "_auto"))

# Combine the new dataframe with the original dataframe
df_combined_area <- df_segManAut_combined %>%
  select(Time, Well, Isolate, pH, Area) %>%
  bind_rows(df_area_auto)

# Make a log area column
df_combined_area <- df_combined_area %>%
  mutate(LogArea = log(Area))
```

When inspecting the graphs, some discrepancies can be observed in the lag phase. To avoid issues we decide to focus on datapoints where log(area) >8, so I make a new dataframe with only those entries.

```
# Make new dataframe with LogArea >8
df_combined_logArea <- df_combined_area %>%
  filter(LogArea > 8)
```

#### Statistically testing if manual and automatic are similar

Start by making a dataframe with "_auto" added to the isolates and area of these moved to the Area column

```
# First making a new dataframe where area is moved to Area and "auto" is added to corresponding isolate

# Step 1: Create new rows with "_auto" added to the Isolate values
new_rows <- df_segManAut_combined %>%
  mutate(Isolate = paste0(Isolate, "_auto"), Area = area) %>%
  select(-area)

# Step 2: Combine new rows with the original dataframe
df_combined <- bind_rows(df_segManAut_combined, new_rows)

# Step 3: removing column area
df_combined_manAut <- df_combined %>%
  select(-area)

#Time has to be > 10 (Edited TIme * 2)
df_combined_manAut$T <- as.numeric(df_combined_manAut$T)
df_combined_manAut <- df_combined_manAut %>%
  filter(T > 18)
```

Then I need to find the values of Time that are behind the highest slope for each isolate:pH combination. I use dpseg for this on log-transformed area. But first, I make subsets of the dataframe for each combination of isolate:pH.

Then I want to create a dataframe that only contains the Area and Time values corresponding to the steepest slope and split this into a subset for each isolate:pH combination.

```
new_subset <- data.frame()

# Loop through each entry in steepest_segments
for (subset_name in names(steepest_segments)) {
  # Extract the time points and isolate from the subset name
  time_points <- steepest_segments[[subset_name]]
  isolate <- gsub("subset_|ph.*", "", subset_name)
  pH_value <- gsub(".*ph", "", subset_name)

  # Determine if the isolate is an _auto isolate
  if (grepl("_auto$", isolate)) {
    # Handle _auto isolates
    filtered_data1 <- df_combined_manAut %>%
      filter(Time %in% time_points, Isolate == isolate, pH == pH_value)
  } else {
    # Handle non-_auto isolates
    filtered_data1 <- df_combined_manAut %>%
      filter(Time %in% time_points, Isolate == isolate, pH == pH_value)
  }

  # Combine the filtered data into the new subset dataframe
  new_subset <- bind_rows(new_subset, filtered_data1)
}
```

```r
# From new_subset, make a new dataframe for each isolate:pH combination (eight new dataframes)
isolates <- c("c005", "c006", "c008", "c009")

# Create an empty list to store the subsets
subsets_timepoints <- list()

#Loop through each combination of Isolate and pH   (OBS: if combining auto and not auto in each subset)
for (isolate in isolates) {
  for (pH_value in pH_values) {
    subset_name <- paste0("subset_", isolate, "ph", gsub("\\.", "", as.character(pH_value)))
    subsets_timepoints[[subset_name]] <- new_subset %>%
      filter(pH == pH_value, str_detect(Isolate, paste0("^", isolate, "(_auto)?$")))
  }
}
```

From the dataframes stored in subsets_timepoints I perform a pairwise comparison using ls.means

```r
# Create an empty list to store the results
results <- list()

# Loop through each isolate and its _auto variant
for (isolate in isolates) {
  for (pH_value in pH_values) {
    # Create a subset for the isolate and its _auto variant at the specific pH level
    subset_data <- new_subset %>%
      filter(str_detect(Isolate, paste0("^", isolate, "(_auto)?$")), pH == pH_value)

    # Fit the linear model
    m.interaction <- lm(Area ~ Time * Isolate, data = subset_data)

    # Perform ANOVA
    anova_result <- anova(m.interaction)

    # Obtain slopes
    m.lst <- lstrends(m.interaction, "Isolate", var = "Time")

    # Compare slopes
    pairs_result <- pairs(m.lst)

    # Summary stats
    summary_result <- summary(m.lst)

    # Store the results
    results[[paste(isolate, pH_value, sep = "_")]] <- list(
      anova = anova_result,
      slopes = m.lst,
      pairs = pairs_result,
      summary = summary_result
    )

    # Plot Area over Time for each subset
    plot(subset_data$Time, subset_data$Area, type = "n", main = paste("Area over Time for", isolate, "a
```

```
    # Add points for the isolate
    isolate_data <- subset_data %>% filter(Isolate == isolate)
    points(isolate_data$Time, isolate_data$Area, col = "black", pch = 16)

    # Add points for the isolate_auto
    isolate_auto_data <- subset_data %>% filter(Isolate == paste0(isolate, "_auto"))
    points(isolate_auto_data$Time, isolate_auto_data$Area, col = "purple", pch = 16)

      # Add linear trendline for the isolate
    isolate_lm <- lm(Area ~ Time, data = isolate_data)
    abline(isolate_lm, col = "black", lwd = 2)

    # Add linear trendline for the isolate_auto
    isolate_auto_lm <- lm(Area ~ Time, data = isolate_auto_data)
    abline(isolate_auto_lm, col = "purple", lwd = 2)

    # Add a legend
    legend("bottomright", legend = c(isolate, paste0(isolate, "_auto")), col = c("black", "purple"), pch
  }
}
```

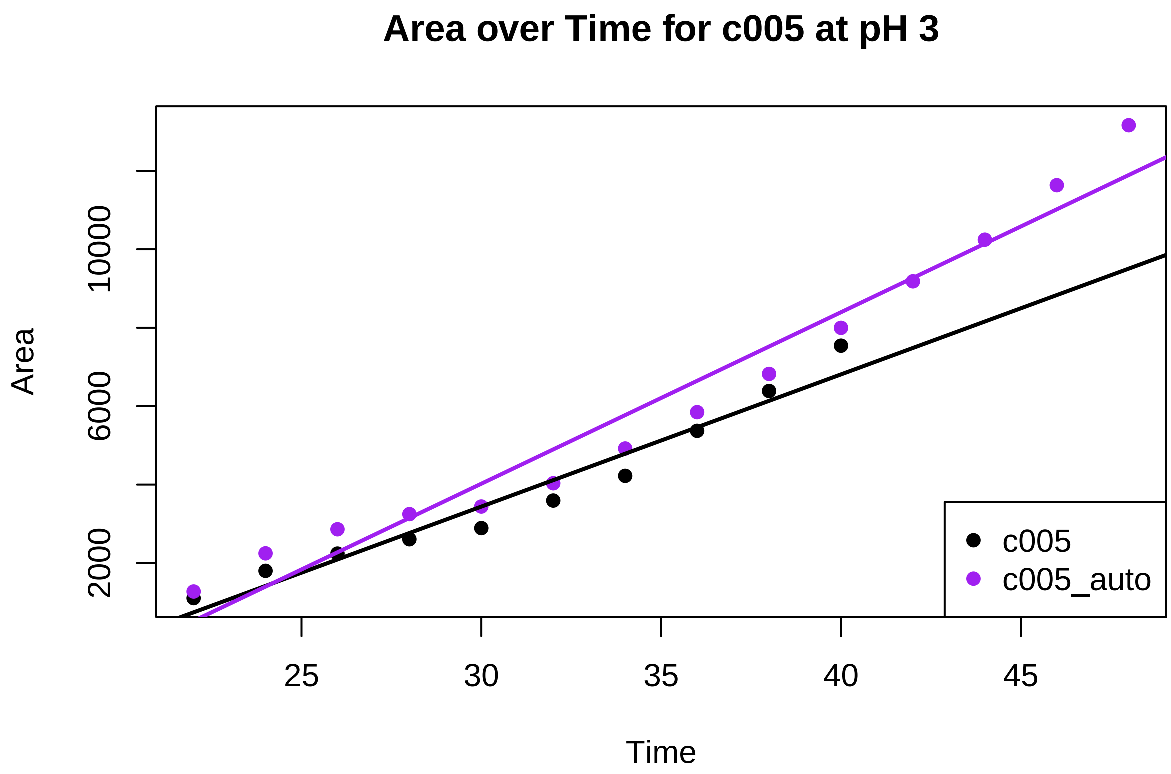

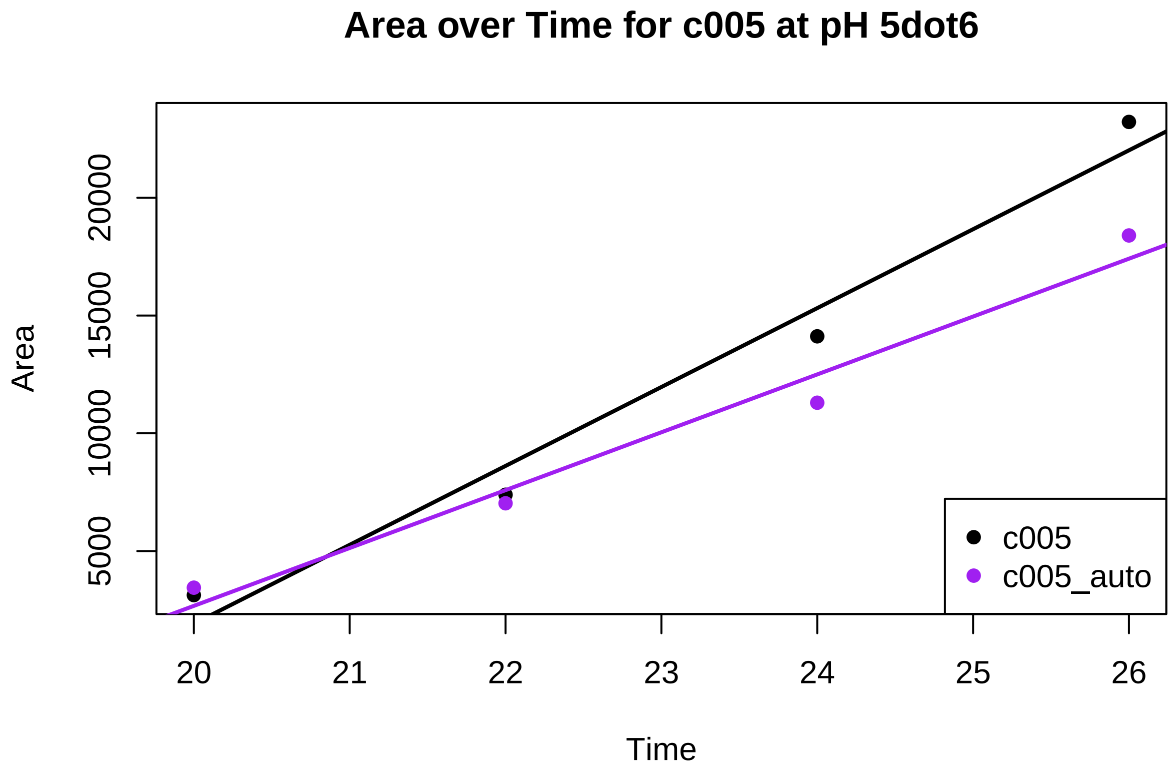
Area over Time for c005 at pH 5dot6
Area
Time
20000
15000
10000
5000
20
21
22
23
24
25
26
c005
c005_auto

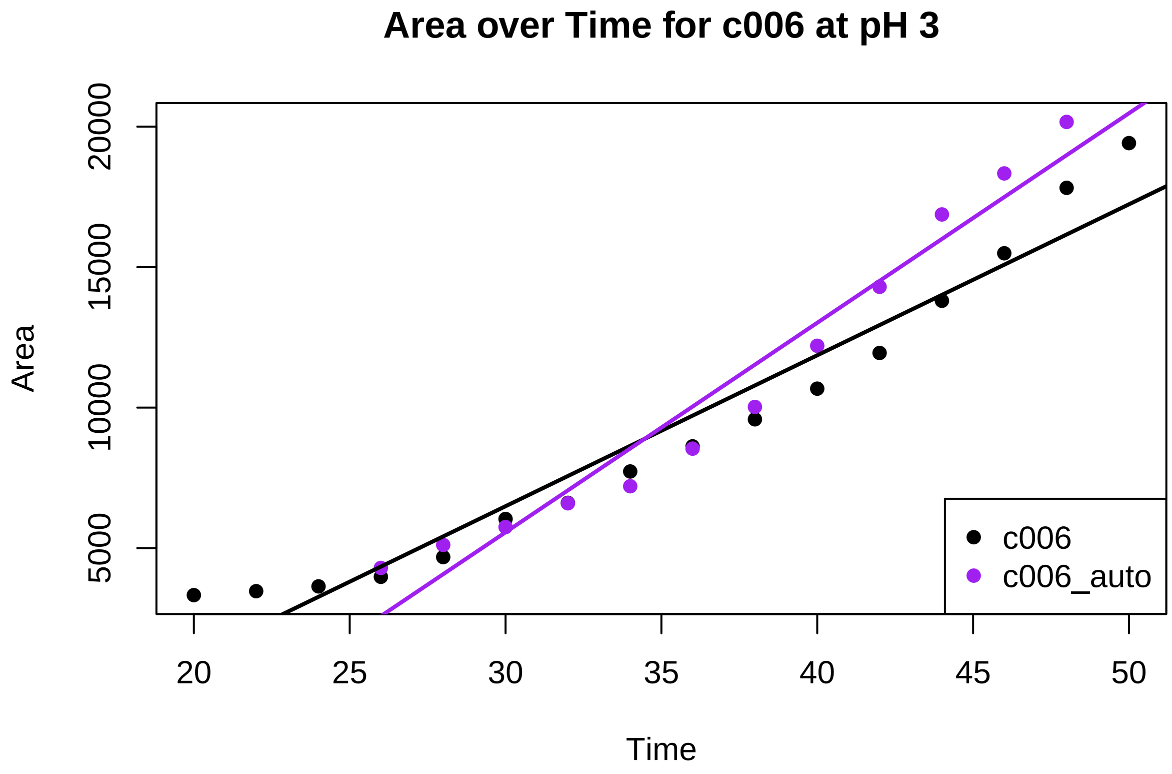

Area over Time for c006 at pH 3
Area
Time
20000
15000
10000
5000
20
25
30
35
40
45
50
c006
c006_auto

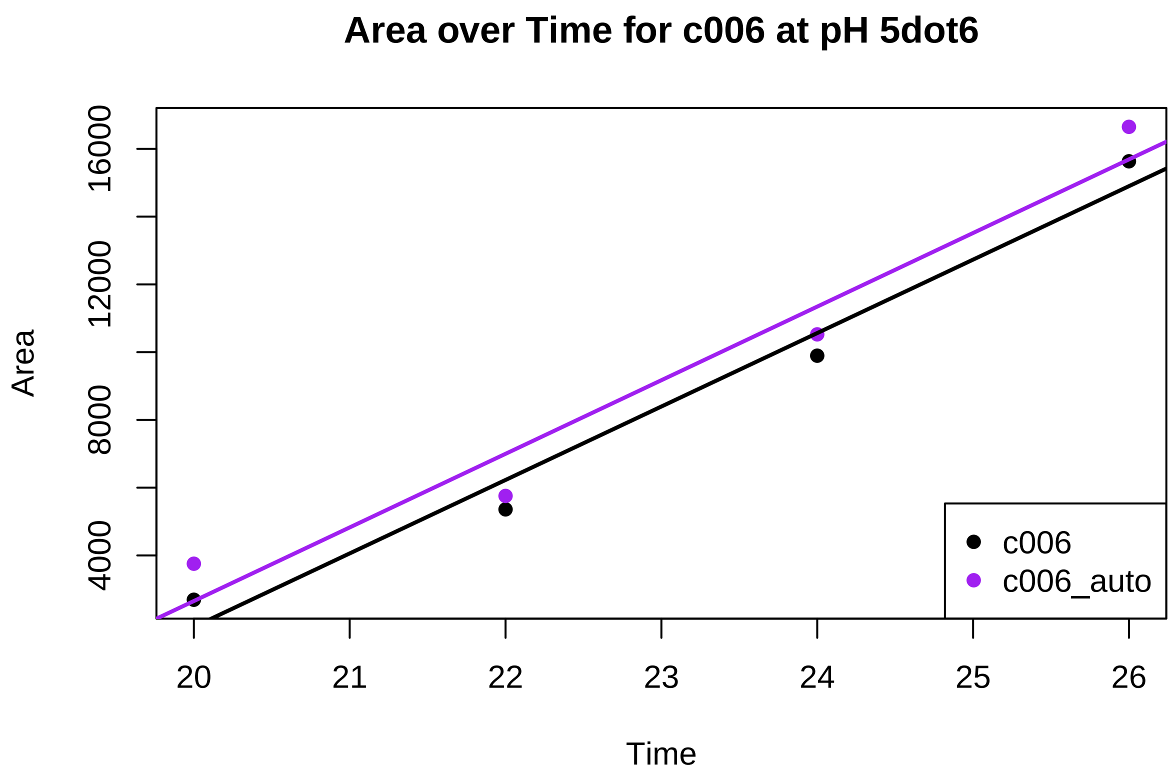
Area over Time for c006 at pH 5dot6
Area
4000
8000
12000
16000
20
21
22
23
24
25
26
Time
c006
c006_auto

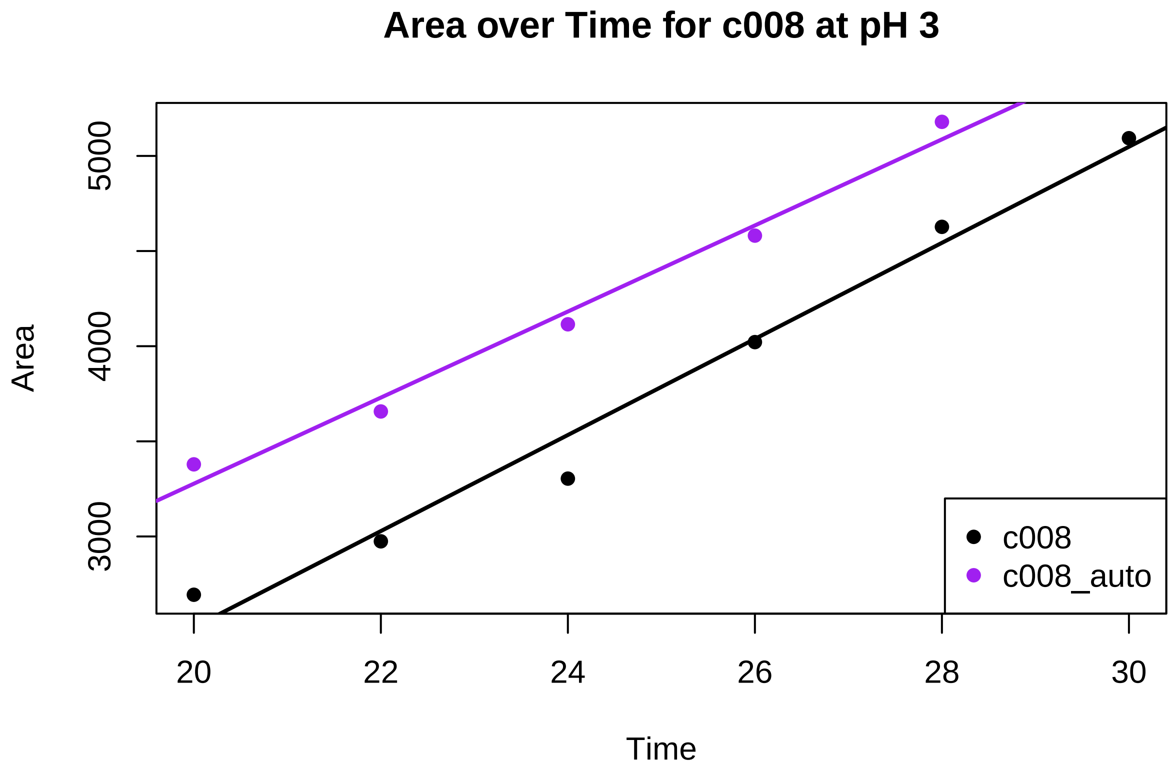

Area over Time for c008 at pH 3
Area
5000
4000
3000
20
22
24
26
28
30
Time
c008
c008_auto

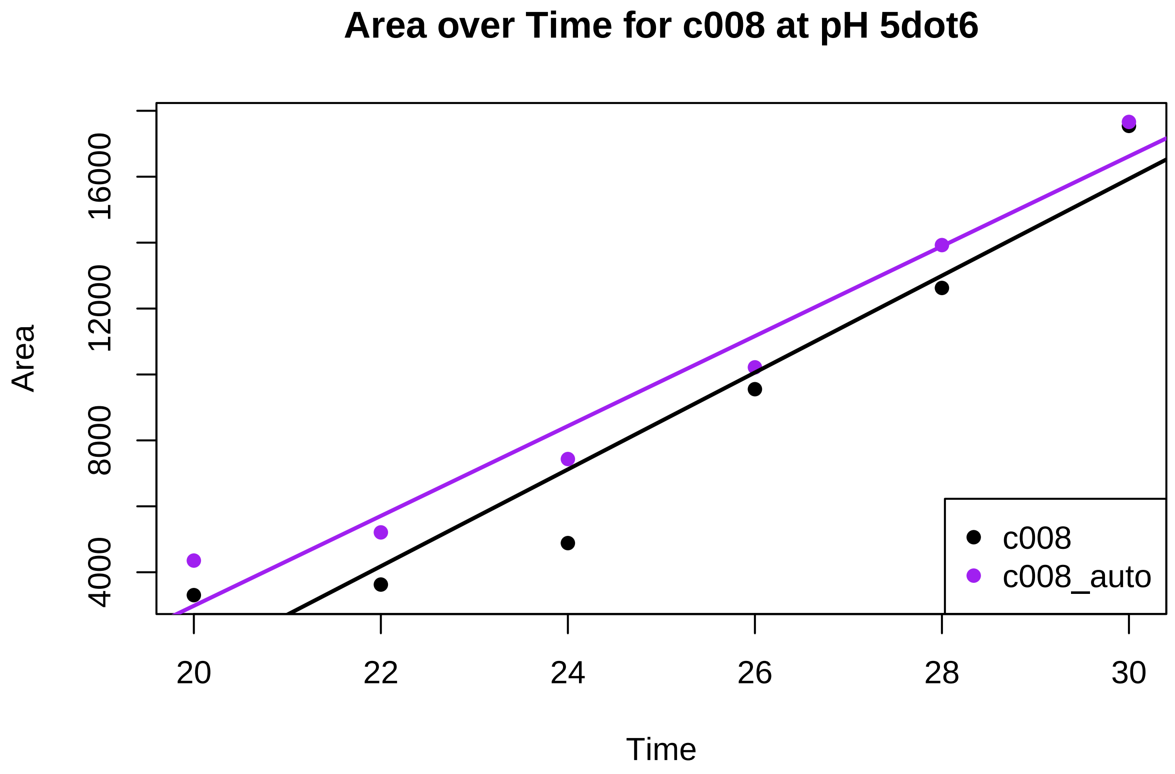
Area over Time for c008 at pH 5dot6
Area
4000
8000
12000
16000
20
22
24
26
28
30
Time
c008
c008_auto

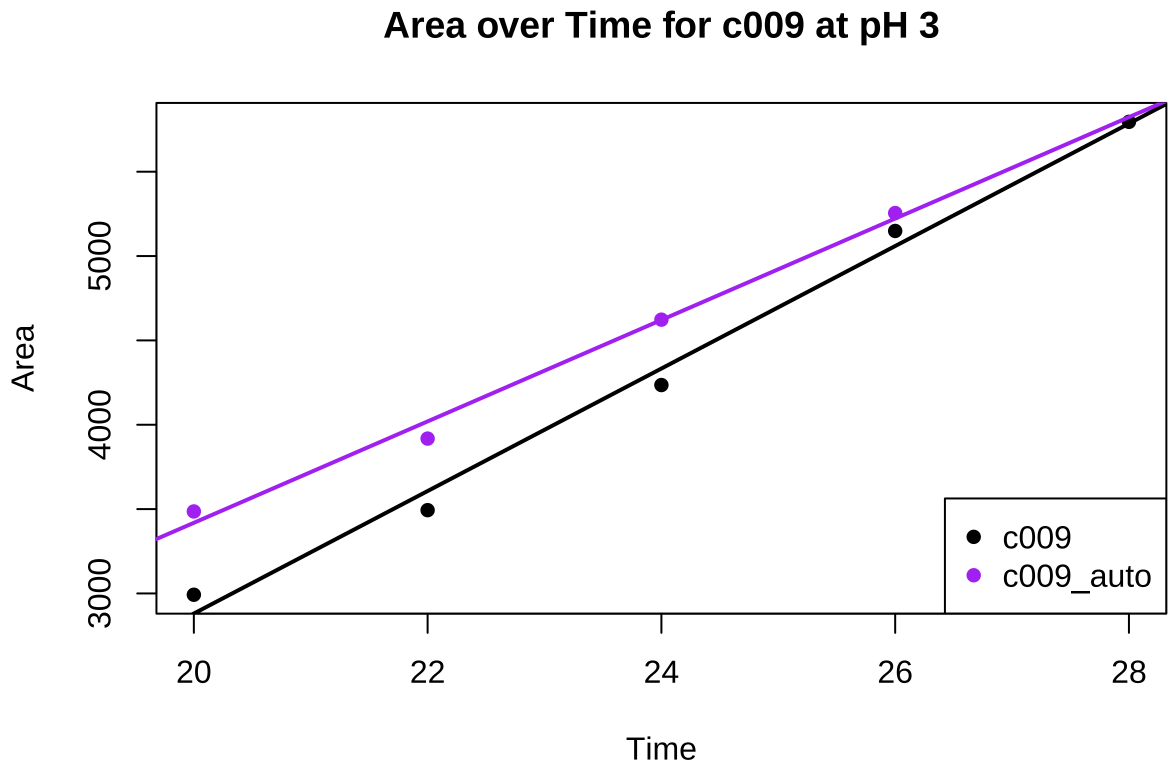

Area over Time for c009 at pH 3
Area
3000
4000
5000
20
22
24
26
28
Time
c009
c009_auto

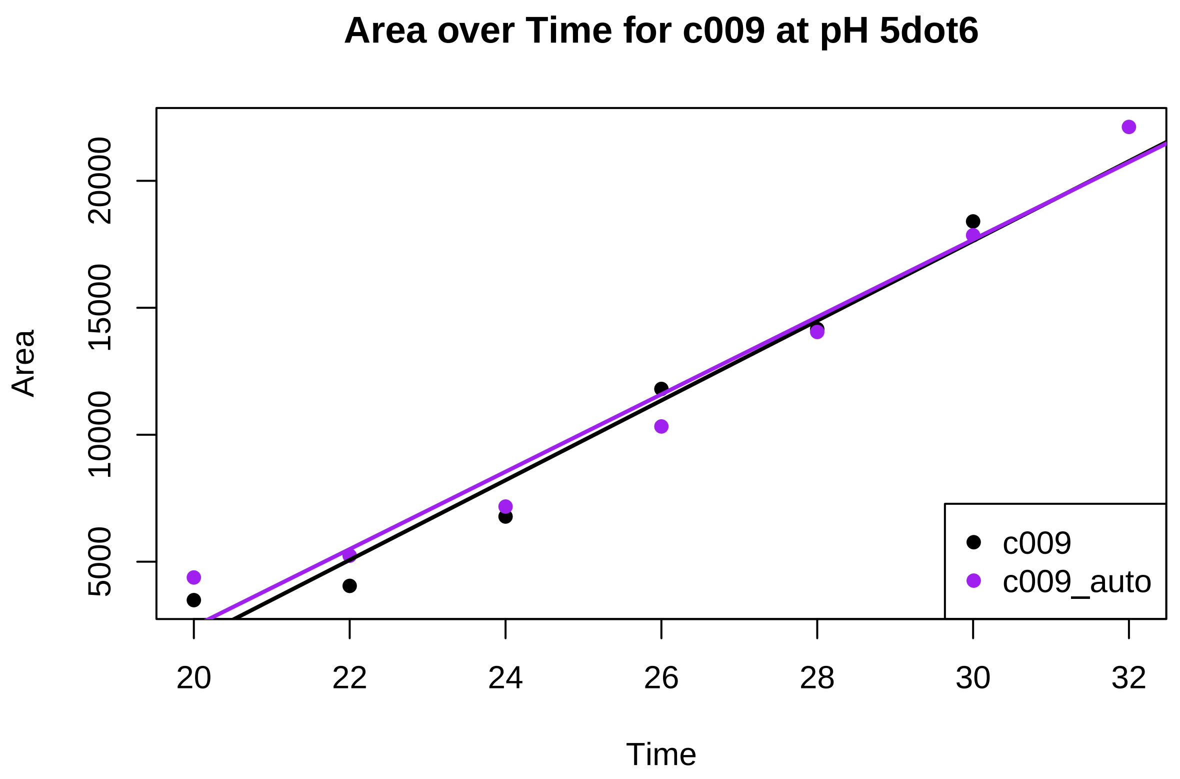

```r
# Loop through each isolate in the results list and print the pairs results and m.lst results
for (result_name in names(results)) {
  cat("\nPairs results for", result_name, ":\n")
  print(results[[result_name]]$pairs)

  cat("\nSlopes (m.lst) results for", result_name, ":\n")
  print(results[[result_name]]$slopes)
}
```

```
##
## Pairs results for c005_3 :
##  contrast         estimate SE df t.ratio p.value
##  c005 - c005_auto     -100 42 20  -2.382  0.0272
##
##
## Slopes (m.lst) results for c005_3 :
##  Isolate   Time.trend   SE df lower.CL upper.CL
##  c005             337 36.0 20      262      412
##  c005_auto        437 21.7 20      392      482
##
## Confidence level used: 0.95
##
## Pairs results for c005_5dot6 :
##  contrast         estimate  SE df t.ratio p.value
##  c005 - c005_auto      893 479  4   1.864  0.1358
##
```

```
##
## Slopes (m.lst) results for c005_5dot6 :
##  Isolate   Time.trend  SE df lower.CL upper.CL
##  c005            3350 339  4     2409     4291
##  c005_auto       2457 339  4     1516     3397
##
## Confidence level used: 0.95
##
## Pairs results for c006_3 :
##  contrast         estimate   SE df t.ratio p.value
##  c006 - c006_auto     -208 61.2 24  -3.406  0.0023
##
##
## Slopes (m.lst) results for c006_3 :
##  Isolate   Time.trend   SE df lower.CL upper.CL
##  c006             537 33.3 24      469      606
##  c006_auto        746 51.3 24      640      852
##
## Confidence level used: 0.95
##
## Pairs results for c006_5dot6 :
##  contrast         estimate  SE df t.ratio p.value
##  c006 - c006_auto    -4.64 410  4  -0.011  0.9915
##
##
## Slopes (m.lst) results for c006_5dot6 :
##  Isolate   Time.trend  SE df lower.CL upper.CL
##  c006            2168 290  4     1363     2973
##  c006_auto       2173 290  4     1367     2978
##
## Confidence level used: 0.95
##
## Pairs results for c008_3 :
##  contrast         estimate   SE df t.ratio p.value
##  c008 - c008_auto     26.3 26.5  7   0.990  0.3550
##
##
## Slopes (m.lst) results for c008_3 :
##  Isolate   Time.trend   SE df lower.CL upper.CL
##  c008             252 16.0  7      215      290
##  c008_auto        226 21.2  7      176      276
##
## Confidence level used: 0.95
##
## Pairs results for c008_5dot6 :
##  contrast         estimate  SE df t.ratio p.value
##  c008 - c008_auto      105 251  8   0.419  0.6859
##
##
## Slopes (m.lst) results for c008_5dot6 :
##  Isolate   Time.trend  SE df lower.CL upper.CL
##  c008            1469 178  8     1060     1879
##  c008_auto       1364 178  8      954     1773
##
```

```
## Confidence level used: 0.95
##
## Pairs results for c009_3 :
##  contrast         estimate   SE df t.ratio p.value
##  c009 - c009_auto     62.5 29.7  5   2.102  0.0895
##
##
## Slopes (m.lst) results for c009_3 :
##  Isolate   Time.trend   SE df lower.CL upper.CL
##  c009             363 17.2  5      319      407
##  c009_auto        301 24.3  5      238      363
##
## Confidence level used: 0.95
##
## Pairs results for c009_5dot6 :
##  contrast         estimate  SE df t.ratio p.value
##  c009 - c009_auto     46.7 203  9   0.230  0.8234
##
##
## Slopes (m.lst) results for c009_5dot6 :
##  Isolate   Time.trend  SE df lower.CL upper.CL
##  c009            1571 159  9     1210     1931
##  c009_auto       1524 126  9     1239     1809
##
## Confidence level used: 0.95
```

We want this pairwise comparison done on only the timepoints overlapping between manual and automatic segmentation. This is not the case for all isolate:pH combinations and I therefore run some of them individually.

(It is repeated for C006:pH 3; c005:pH3; C008:pH3 and C009:pH5.6 )

###### Using same datapoints for C006 ph 3:

```
# Obtain slopes
m.interaction$coefficients
```

```
##           (Intercept)                  Time      Isolatec006_auto
##           -12277.9714              595.2839            -4529.9039
## Time:Isolatec006_auto
##              150.3996
```

```
m.lst <- lstrends(m.interaction, "Isolate", var="Time")

m.lst
```

```
##  Isolate   Time.trend   SE df lower.CL upper.CL
##  c006             595 41.6 20      508      682
##  c006_auto        746 41.6 20      659      833
##
## Confidence level used: 0.95
```

```
# Compare slopes
#significant difference?
pairs(m.lst)
```

```
##  contrast         estimate   SE df t.ratio p.value
##  c006 - c006_auto     -150 58.9 20  -2.554  0.0189
```

```
# summary stats
summary(m.lst)
```

```
##  Isolate   Time.trend   SE df lower.CL upper.CL
##  c006             595 41.6 20      508      682
##  c006_auto        746 41.6 20      659      833
##
## Confidence level used: 0.95
```

```
# Define the isolate and pH value you want to plot
isolate <- "c006"
pH_value <- 3

# Create a subset for the isolate and its _auto variant at the specific pH level
subset_data <- subset_c006_ph3 %>%
  filter(str_detect(Isolate, paste0("^", isolate, "(_auto)?$")), pH == pH_value)

# Plot Area over Time for the subset
plot(subset_data$Time, subset_data$Area, type = "n", main = paste("Area over Time for", isolate, "at pH

# Add points for the isolate
isolate_data <- subset_data %>% filter(Isolate == isolate)
points(isolate_data$Time, isolate_data$Area, col = "black", pch = 16)

# Add points for the isolate_auto
isolate_auto_data <- subset_data %>% filter(Isolate == paste0(isolate, "_auto"))
points(isolate_auto_data$Time, isolate_auto_data$Area, col = "purple", pch = 16)

# Add linear trendline for the isolate
isolate_lm <- lm(Area ~ Time, data = isolate_data)
abline(isolate_lm, col = "black", lwd = 2)

# Add linear trendline for the isolate_auto
isolate_auto_lm <- lm(Area ~ Time, data = isolate_auto_data)
abline(isolate_auto_lm, col = "purple", lwd = 2)

# Add a legend
legend("bottomright", legend = c(isolate, paste0(isolate, "_auto")), col = c("black", "purple"), pch = 
```

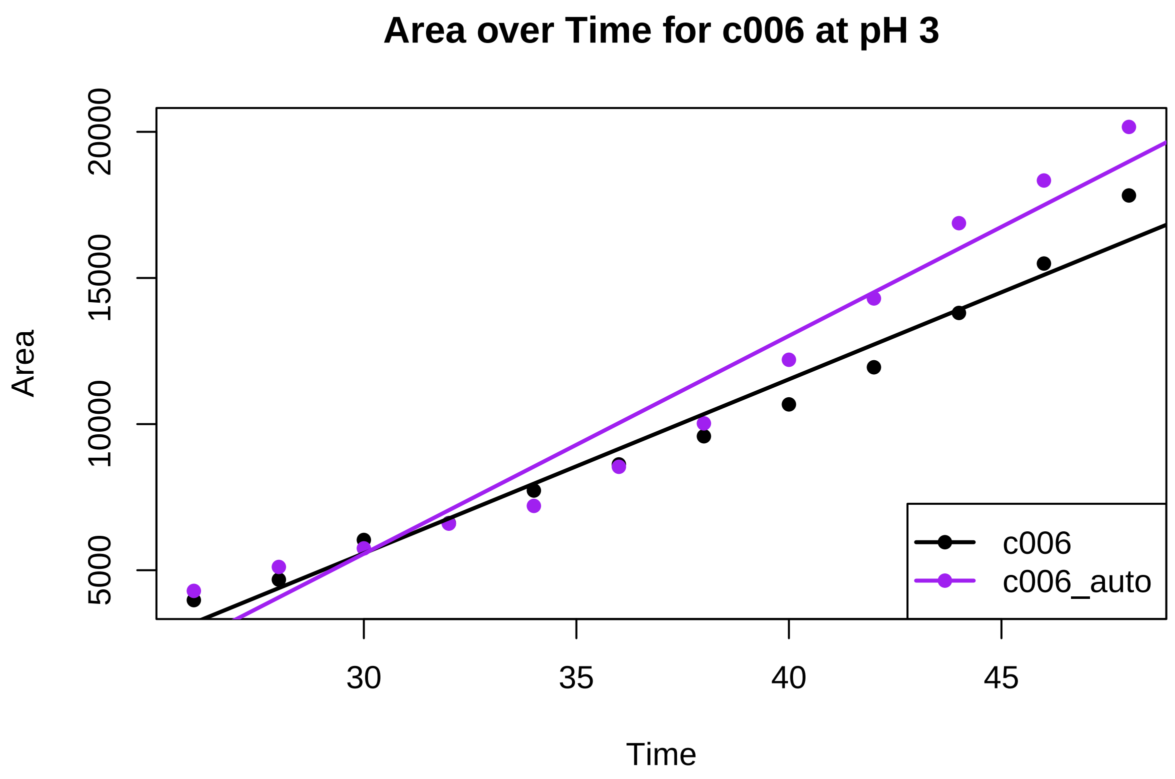

**Using same timepoints for C005 at pH3:**

```
# Obtain slopes
m.interaction$coefficients
```

```
##           (Intercept)                  Time      Isolatec005_auto
##          -6673.373404            337.077620            320.270373
## Time:Isolatec005_auto
##              5.570865
```

```
m.lst <- lstrends(m.interaction, "Isolate", var="Time")

m.lst
```

```
##  Isolate   Time.trend   SE df lower.CL upper.CL
##  c005             337 24.8 16      285      390
##  c005_auto        343 24.8 16      290      395
##
## Confidence level used: 0.95
```

```
# Compare slopes
#significant difference?
pairs(m.lst)
```

```
##  contrast         estimate SE df t.ratio p.value
##  c005 - c005_auto    -5.57 35 16  -0.159  0.8756
```

```r
# summary stats
summary(m.lst)
```

```
##  Isolate   Time.trend   SE df lower.CL upper.CL
##  c005             337 24.8 16      285      390
##  c005_auto        343 24.8 16      290      395
##
## Confidence level used: 0.95
```

```r
# Define the isolate and pH value you want to plot
isolate <- "c005"
pH_value <- 3

# Create a subset for the isolate and its _auto variant at the specific pH level
subset_data <- subset_c005_ph3 %>%
  filter(str_detect(Isolate, paste0("^", isolate, "(_auto)?$")), pH == pH_value)

# Plot Area over Time for the subset
plot(subset_data$Time, subset_data$Area, type = "n", main = paste("Area over Time for", isolate, "at pH

# Add points for the isolate
isolate_data <- subset_data %>% filter(Isolate == isolate)
points(isolate_data$Time, isolate_data$Area, col = "black", pch = 16)

# Add points for the isolate_auto
isolate_auto_data <- subset_data %>% filter(Isolate == paste0(isolate, "_auto"))
points(isolate_auto_data$Time, isolate_auto_data$Area, col = "purple", pch = 16)

# Add linear trendline for the isolate
isolate_lm <- lm(Area ~ Time, data = isolate_data)
abline(isolate_lm, col = "black", lwd = 2)

# Add linear trendline for the isolate_auto
isolate_auto_lm <- lm(Area ~ Time, data = isolate_auto_data)
abline(isolate_auto_lm, col = "purple", lwd = 2)

# Add a legend
legend("bottomright", legend = c(isolate, paste0(isolate, "_auto")), col = c("black", "purple"), pch =
```

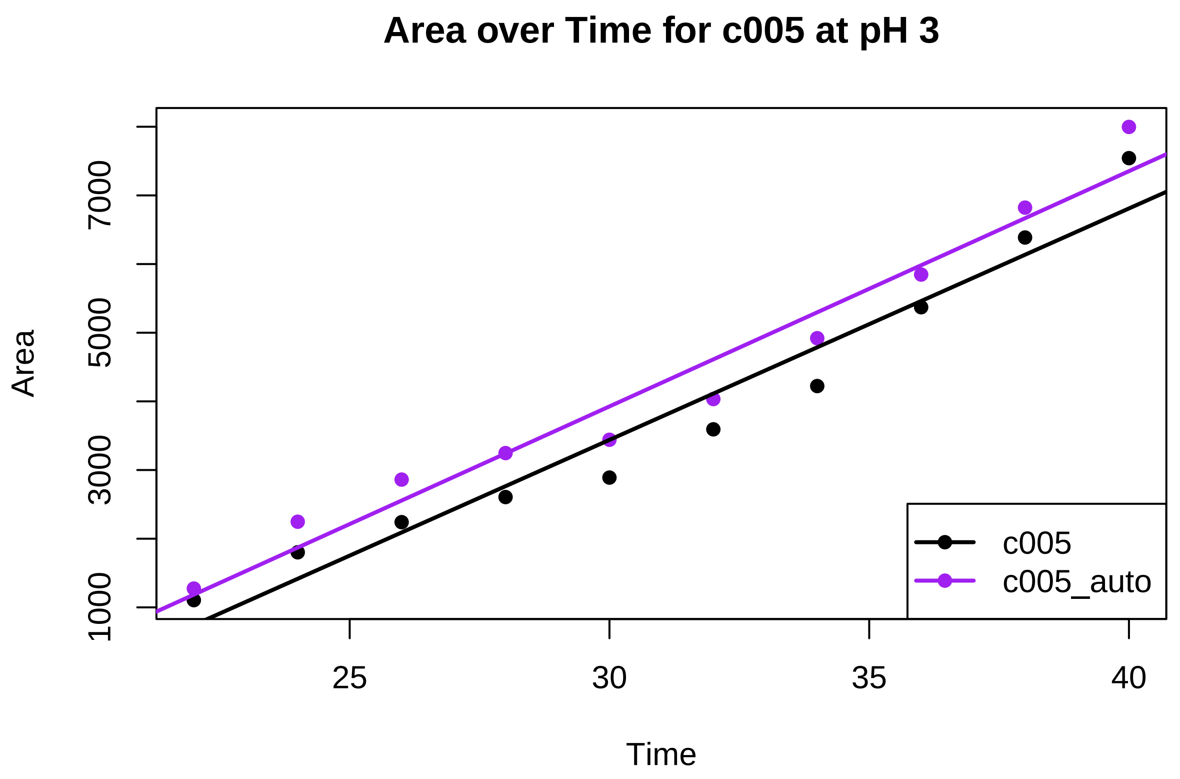

**Using same timepoints for C008 pH 3 :**

```
# Obtain slopes
m.interaction$coefficients
```

```
##           (Intercept)                 Time      Isolatec008_auto
##           -2372.47074            245.69323            1125.87074
## Time:Isolatec008_auto
##             -19.49323
```

```
m.lst <- lstrends(m.interaction, "Isolate", var="Time")

m.lst
```

```
##  Isolate   Time.trend   SE df lower.CL upper.CL
##  c008             246 22.5  6      191      301
##  c008_auto        226 22.5  6      171      281
##
## Confidence level used: 0.95
```

```
# Compare slopes
#significant difference?
pairs(m.lst)
```

```
##  contrast          estimate   SE df t.ratio p.value
##  c008 - c008_auto     19.5 31.8  6   0.614  0.5619
```

```
# summary stats
summary(m.lst)
```

```
##  Isolate   Time.trend   SE df lower.CL upper.CL
##  c008             246 22.5  6      191      301
##  c008_auto        226 22.5  6      171      281
##
## Confidence level used: 0.95
```

```
# Define the isolate and pH value you want to plot
isolate <- "c008"
pH_value <- 3

# Create a subset for the isolate and its _auto variant at the specific pH level
subset_data <- subset_c008_ph3 %>%
  filter(str_detect(Isolate, paste0("^", isolate, "(_auto)?$")), pH == pH_value)

# Plot Area over Time for the subset
plot(subset_data$Time, subset_data$Area, type = "n", main = paste("Area over Time for", isolate, "at pH

# Add points for the isolate
isolate_data <- subset_data %>% filter(Isolate == isolate)
points(isolate_data$Time, isolate_data$Area, col = "black", pch = 16)

# Add points for the isolate_auto
isolate_auto_data <- subset_data %>% filter(Isolate == paste0(isolate, "_auto"))
points(isolate_auto_data$Time, isolate_auto_data$Area, col = "purple", pch = 16)

# Add linear trendline for the isolate
isolate_lm <- lm(Area ~ Time, data = isolate_data)
abline(isolate_lm, col = "black", lwd = 2)

# Add linear trendline for the isolate_auto
isolate_auto_lm <- lm(Area ~ Time, data = isolate_auto_data)
abline(isolate_auto_lm, col = "purple", lwd = 2)

# Add a legend
legend("bottomright", legend = c(isolate, paste0(isolate, "_auto")), col = c("black", "purple"), pch =
```

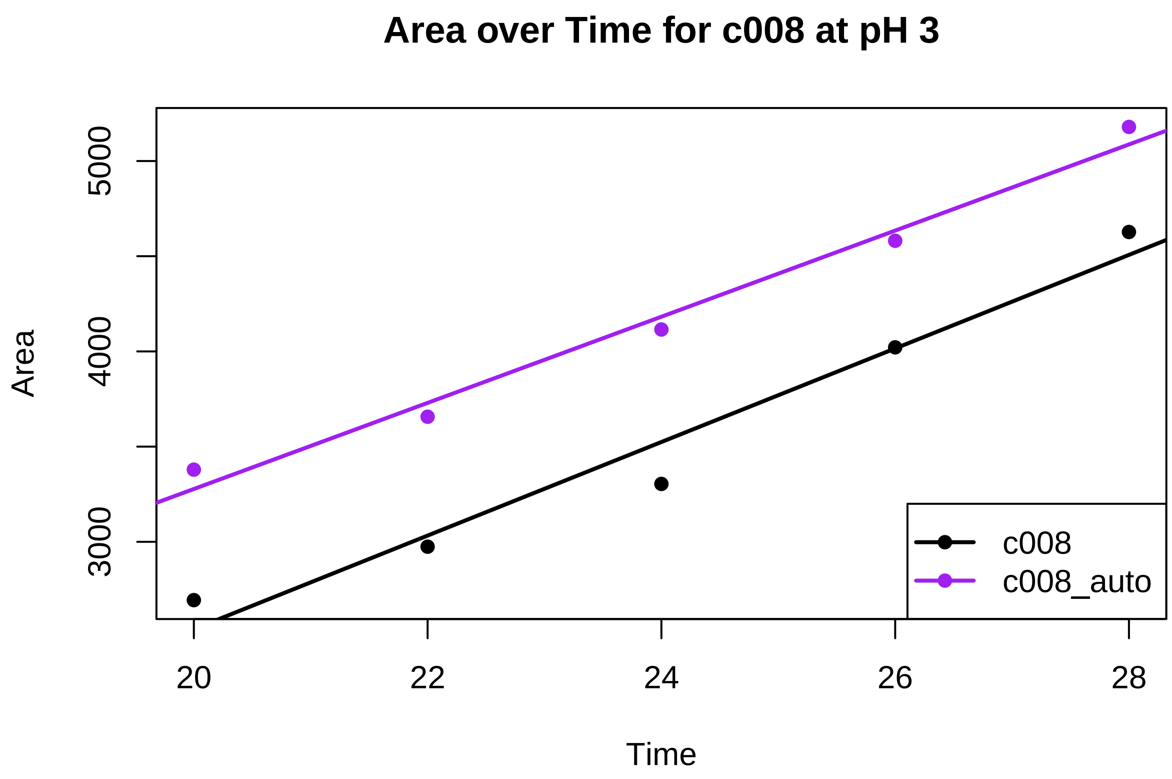

**Using same timepoints for C009 ph 5.6 :**

```
# Obtain slopes
m.interaction$coefficients
```

```
##           (Intercept)                  Time       Isolatec009_auto
##           -29482.5940             1570.5235              4691.2130
## Time:Isolatec009_auto
##             -185.3949
```

```
m.lst <- lstrends(m.interaction, "Isolate", var="Time")

m.lst
```

```
##  Isolate   Time.trend  SE df lower.CL upper.CL
##  c009            1571 149  8     1227     1914
##  c009_auto       1385 149  8     1042     1728
##
## Confidence level used: 0.95
```

```
# Compare slopes
#significant difference?
pairs(m.lst)
```

```
##  contrast          estimate  SE df t.ratio p.value
##  c009 - c009_auto      185 211  8   0.881  0.4043
```

```
# summary stats
summary(m.lst)
```

```
##  Isolate   Time.trend  SE df lower.CL upper.CL
##  c009            1571 149  8     1227     1914
##  c009_auto       1385 149  8     1042     1728
##
## Confidence level used: 0.95
```

```
# Define the isolate and pH value you want to plot
isolate <- "c009"
pH_value <- "5dot6"

# Create a subset for the isolate and its _auto variant at the specific pH level
subset_data <- subset_c009_ph5dot6 %>%
  filter(str_detect(Isolate, paste0("^", isolate, "(_auto)?$")), pH == pH_value)

# Plot Area over Time for the subset
plot(subset_data$Time, subset_data$Area, type = "n", main = paste("Area over Time for", isolate, "at pH

# Add points for the isolate
isolate_data <- subset_data %>% filter(Isolate == isolate)
points(isolate_data$Time, isolate_data$Area, col = "black", pch = 16)

# Add points for the isolate_auto
isolate_auto_data <- subset_data %>% filter(Isolate == paste0(isolate, "_auto"))
points(isolate_auto_data$Time, isolate_auto_data$Area, col = "purple", pch = 16)

# Add linear trendline for the isolate
isolate_lm <- lm(Area ~ Time, data = isolate_data)
abline(isolate_lm, col = "black", lwd = 2)

# Add linear trendline for the isolate_auto
isolate_auto_lm <- lm(Area ~ Time, data = isolate_auto_data)
abline(isolate_auto_lm, col = "purple", lwd = 2)

# Add a legend
legend("bottomright", legend = c(isolate, paste0(isolate, "_auto")), col = c("black", "purple"), pch = 
```

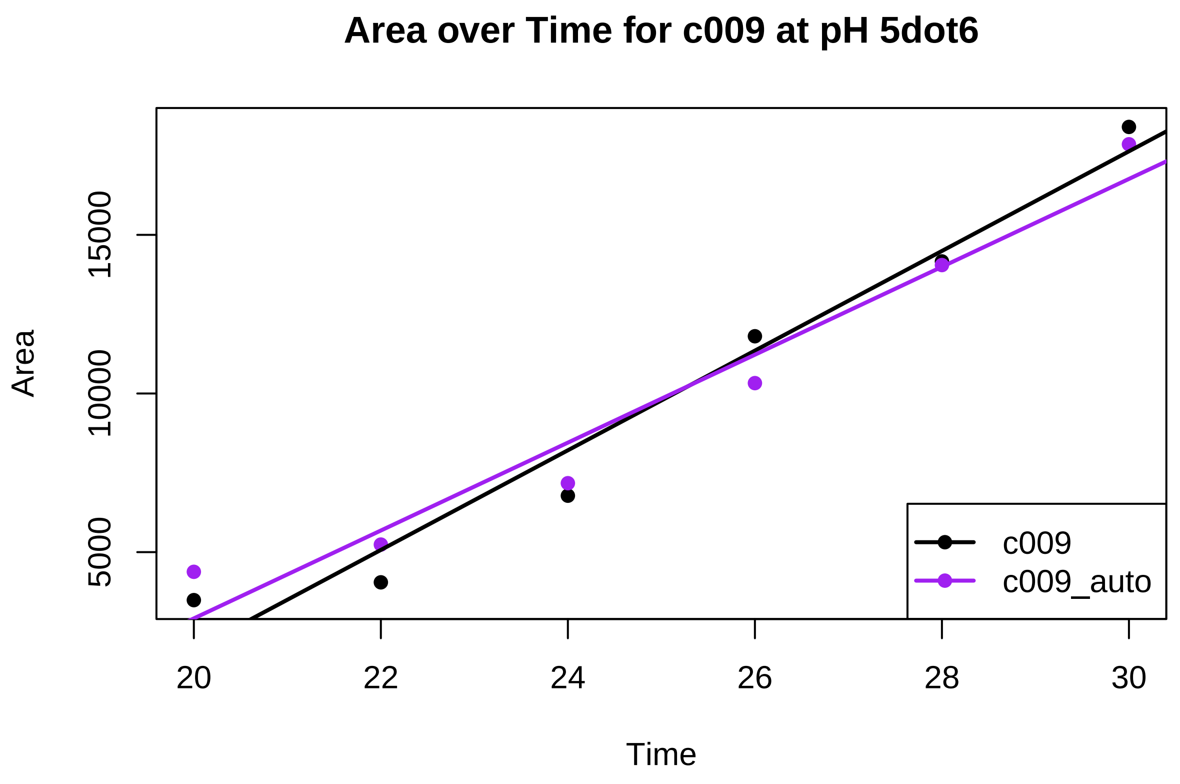

**Multiple comparison correction with Benjamini & Hochberg**

I perform multiple comparison correction with Benjamini & Hcohberg method. I do it on the p-values I have found when only using the overlapping timepoints between manual and auto. I write these manually into a numeric vectorp (in this order: C005 ph3, c006 ph 3, c008 ph 3, c009 ph 3, c005 ph 5.6, c006 ph 5.6, c008 ph 5.6, c009 ph 5.6) and then use p.adjust:

```
# Manually write the p-values in numeric vector p:
p <- c(0.8756, 0.0189, 0.5619, 0.0895, 0.1358, 0.9915, 0.6859, 0.4043)

# Applying p.adjust with Benjamini & Hochberg:
BH_adjusted <- p.adjust(p, method = "BH", n = length(p))

# Print the adjusted p-values for each method
print("Benjamini & Hochberg adjusted p-values:")
```

```
## [1] "Benjamini & Hochberg adjusted p-values:"
```

```
print(BH_adjusted)
```

```
## [1] 0.9915000 0.1512000 0.8990400 0.3580000 0.3621333 0.9915000 0.9145333
## [8] 0.8086000
```

### Illustration of method on full dataset of pH 3 and pH 5.6 (Test set 2)

We can now argue that the deep learning model is working. In the following, we show how it can be used to segment a lot of images fast, by using it on all the replicates in the experiment with the combinations of isolates C005, C006, C008, C009 and pH 3 and 5.6.

#### Tidying dataframe

I use the dataframe df_auto that we already loaded previously, when comparing manual and automated measurements. But now instead of only looking at one well per isolate:pH combination, we will use the full dataset of pH 3 and pH 5.6 to demonstrate the method.

First I tidy the dataframe and make sure that log(Area) > 8 to make sure that discreprancies in the lag phase will not cause problems. I also make sure that Time is not above 84 (bug from the imaging system) and that the dataframe contains the column ReplicateNumber.

```
# I also make a column called ReplicateNumber based on the Well column but only keeping the number in t
df_auto_repl <- df_auto %>%
  mutate(ReplicateNumber = substr(Well, 2, 2)         )

df_auto_repl <- df_auto_repl %>%
  filter(log(area) > 8)

#somehow a few T>42 has found it's way in here, so I delete these (edit to T * 2):
df_auto_repl <- df_auto_repl %>%
  filter(Time <= 84)

# Similarly there are pH levels other than 3 and 5dot6 so I remove these
df_auto_repl <- df_auto_repl %>%
  filter(pH %in% c("3", "5dot6"))
```

#### Fitting growth model and extracting maximum growth rate

Then use the function fit_easylinear from the package growthrate in a loop to do it for each replicate of each isolate:pH combination.

Extracting maximum growthrate for each combination of isolate:pH.

```
result_list <- list()

# Loop through all replicates
for (i in 1:48) {
  dat <- splitted.data[[i]]

  # Fit the model
  fit <- fit_easylinear(dat$Time, dat$area)

  # Print coefficients
  mumax <- coef(fit)
  # Print R-squared
  Rsqr <- rsquared(fit)
```

```
# Put all the results into a list for printing
  result_list[[i]] <- list(mumax = mumax, Rsqr = Rsqr)
}

df_mumaxRsqr <- data.frame(result_list)

# Extract the mumax values and R2 from the third row.
selected_row <- df_mumaxRsqr[3, ]
original_row <- unlist(selected_row)

# Split the original row into pairs and create a new data frame
new_df <- data.frame(matrix(original_row, ncol = 2, byrow = TRUE))

# Rename the columns
colnames(new_df) <- c("mumax", "R2")

#Bind this to combination of Isolate, pH and Well
tibble_names <- names(splitted.data)
new_df$TibbleNames <- tibble_names[1:nrow(new_df)]

#Sort by the 'TibbleNames' column
sorted_df_091224 <- new_df %>% arrange(TibbleNames)

# Print the sorted dataframe
print(sorted_df_091224)
```

```
##         mumax        R2  TibbleNames
## 1  0.08521892 0.9984344     c005:3:1
## 2  0.09337849 0.9961699     c005:3:2
## 3  0.08422668 0.9973284     c005:3:3
## 4  0.08506610 0.9981777     c005:3:4
## 5  0.08636282 0.9971403     c005:3:5
## 6  0.07591028 0.9975960     c005:3:6
## 7  0.24711318 0.9811517 c005:5dot6:1
## 8  0.27945641 0.9852491 c005:5dot6:2
## 9  0.20213552 0.9991251 c005:5dot6:3
## 10 0.29587361 0.9937680 c005:5dot6:4
## 11 0.25727613 0.9907904 c005:5dot6:5
## 12 0.21164673 0.9842749 c005:5dot6:6
## 13 0.08573199 0.9993349     c006:3:1
## 14 0.05715124 0.9992337     c006:3:2
## 15 0.06627527 0.9952897     c006:3:3
## 16 0.05587257 0.9981708     c006:3:4
## 17 0.06553082 0.9981559     c006:3:5
## 18 0.06821331 0.9993826     c006:3:6
## 19 0.23661866 0.9925077 c006:5dot6:1
## 20 0.21117210 0.9594100 c006:5dot6:2
## 21 0.28673988 0.9378155 c006:5dot6:3
## 22 0.19289913 0.9466997 c006:5dot6:4
## 23 0.20943852 0.9651249 c006:5dot6:5
## 24 0.17725060 0.9972552 c006:5dot6:6
```

```
## 25 0.05246647 0.9960405     c008:3:1
## 26 0.05589995 0.9949606     c008:3:2
## 27 0.05721183 0.9989676     c008:3:3
## 28 0.05679994 0.9923476     c008:3:4
## 29 0.05862983 0.9899753     c008:3:5
## 30 0.06887624 0.9970891     c008:3:6
## 31 0.14670693 0.9942008 c008:5dot6:1
## 32 0.13841695 0.9970213 c008:5dot6:2
## 33 0.12610335 0.9997890 c008:5dot6:3
## 34 0.10474177 0.9905291 c008:5dot6:4
## 35 0.11216762 0.9949445 c008:5dot6:5
## 36 0.12043107 0.9968323 c008:5dot6:6
## 37 0.06540040 0.9930998     c009:3:1
## 38 0.06738422 0.9917820     c009:3:2
## 39 0.07297986 0.9934091     c009:3:3
## 40 0.07094008 0.9950929     c009:3:4
## 41 0.07204888 0.9966164     c009:3:5
## 42 0.05924734 0.9969927     c009:3:6
## 43 0.14786047 0.9927948 c009:5dot6:1
## 44 0.12946781 0.9947570 c009:5dot6:2
## 45 0.16633132 0.9782973 c009:5dot6:3
## 46 0.15173942 0.9980720 c009:5dot6:4
## 47 0.14025969 0.9966800 c009:5dot6:5
## 48 0.13062747 0.9979896 c009:5dot6:6
```

```
write.csv(sorted_df_091224, "mumax-rsquared_Dataset2_091224.csv", row.names = FALSE)
```

##### Plotting maximum growth rate for all replicates

Plotting all replicates of each combination of isolate and pH as jitterplots.

```
# Splitting replicate from isolate and pH in dataframe
sorted_df_091224 <- sorted_df_091224 %>%
  separate(TibbleNames, into = c("Category", "replicate"), sep = "(?<=:)(?=[^:]+$)")

# Making category column for coloring
sorted_df_091224 <- sorted_df_091224 %>%
  mutate(new_category = factor(Category, labels = LETTERS[1:length(unique(Category))]))

# Plotting, coloring per isolate and inserting a mean
jitter <- ggplot(sorted_df_091224, aes(x = Category, y = mumax, color = new_category)) +
  geom_jitter(width = 0.2, height = 0, alpha = 0.6) +
  stat_summary(fun = mean, geom = "point", shape = 18, size = 3, color = "black") +
  theme_minimal() +
  labs(title = "Maximum growth rate for each Isolate:pH combination", x = "Isolate:pH combination", y =
  scale_color_manual(values = c("A" = "darkred", "B" = "lightcoral",
                                "C" = "darkorange", "D" = "lightsalmon",
                                "E" = "darkgreen", "F" = "lightgreen",
                                "G" = "darkblue", "H" = "lightblue")) +
  theme(axis.text.x = element_text(angle = 310, hjust = 0))

plot(jitter)
```

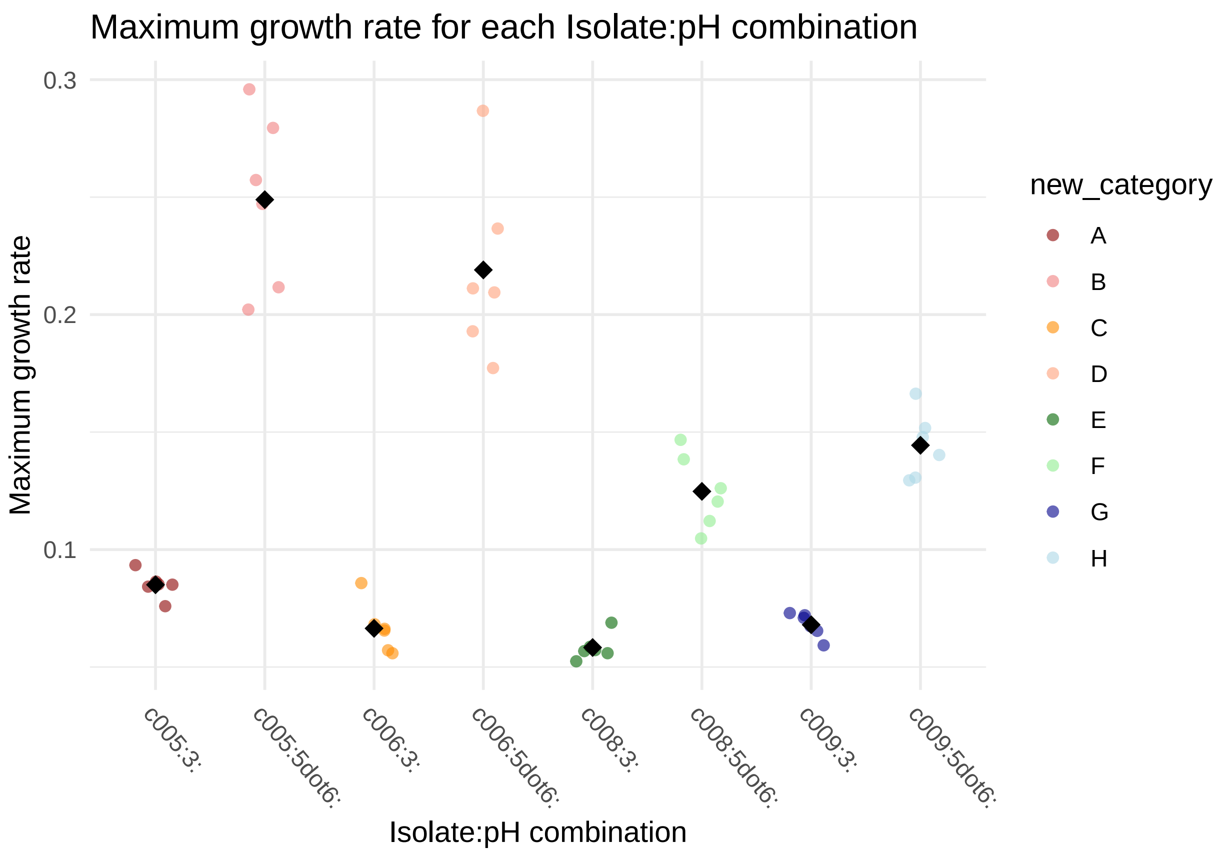

And printing the means of the replicates:

```
# Calculating means for each category
means <- sorted_df_091224 %>%
  group_by(Category) %>%
  summarise(mean_mumax = mean(mumax, na.rm = TRUE))

# Printing the means
print(means)
```

```
## # A tibble: 8 x 2
##   Category    mean_mumax
##   <chr>            <dbl>
## 1 c005:3:         0.0850
## 2 c005:5dot6:     0.249
## 3 c006:3:         0.0665
## 4 c006:5dot6:     0.219
## 5 c008:3:         0.0583
## 6 c008:5dot6:     0.125
## 7 c009:3:         0.0680
## 8 c009:5dot6:     0.144
```

##### Statistical analysis (inter- and intravariation)

We want to look at the variance within and between isolate:pH combinations (inter- and intravariation).

I run ANOVA and perform Tukey's to compare the categories (isolate:pH combinations) pairwise:

```r
# Fit the model
model <- aov(mumax ~ Category, data = sorted_df_091224)

# Get the ANOVA table
anova_result <- summary(model)
print(anova_result)
```

```
##             Df  Sum Sq Mean Sq F value Pr(>F)
## Category     7 0.22358 0.03194   72.98 <2e-16 ***
## Residuals   40 0.01751 0.00044
## ---
## Signif. codes:  0 '***' 0.001 '**' 0.01 '*' 0.05 '.' 0.1 ' ' 1
```

Tukey's HSD test to make pairwise comparison between categories (isolate:pH combinations):

```r
# Perform Tukey's HSD test
tukey_result <- TukeyHSD(model)

# View the results
print(tukey_result)
```

```
##   Tukey multiple comparisons of means
##     95% family-wise confidence level
##
## Fit: aov(formula = mumax ~ Category, data = sorted_df_091224)
##
## $Category
##                                 diff          lwr          upr     p adj
## c005:5dot6:-c005:3:      0.163889714  0.125280327  0.202499100 0.0000000
## c006:3:-c005:3:         -0.018564680 -0.057174066  0.020044706 0.7830431
## c006:5dot6:-c005:3:      0.133992600  0.095383214  0.172601986 0.0000000
## c008:3:-c005:3:         -0.026713171 -0.065322557  0.011896215 0.3667742
## c008:5dot6:-c005:3:      0.039734067  0.001124681  0.078343453 0.0397626
## c009:3:-c005:3:         -0.017027085 -0.055636472  0.021582301 0.8474502
## c009:5dot6:-c005:3:      0.059353816  0.020744430  0.097963202 0.0003842
## c006:3:-c005:5dot6:     -0.182454394 -0.221063780 -0.143845008 0.0000000
## c006:5dot6:-c005:5dot6: -0.029897113 -0.068506499  0.008712273 0.2353352
## c008:3:-c005:5dot6:     -0.190602885 -0.229212271 -0.151993499 0.0000000
## c008:5dot6:-c005:5dot6: -0.124155647 -0.162765033 -0.085546261 0.0000000
## c009:3:-c005:5dot6:     -0.180916799 -0.219526185 -0.142307413 0.0000000
## c009:5dot6:-c005:5dot6: -0.104535898 -0.143145284 -0.065926512 0.0000000
## c006:5dot6:-c006:3:      0.152557280  0.113947894  0.191166666 0.0000000
## c008:3:-c006:3:         -0.008148491 -0.046757877  0.030460895 0.9972423
## c008:5dot6:-c006:3:      0.058298747  0.019689361  0.096908133 0.0005033
## c009:3:-c006:3:          0.001537595 -0.037071791  0.040146981 1.0000000
## c009:5dot6:-c006:3:      0.077918496  0.039309110  0.116527882 0.0000029
## c008:3:-c006:5dot6:     -0.160705772 -0.199315158 -0.122096385 0.0000000
## c008:5dot6:-c006:5dot6: -0.094258534 -0.132867920 -0.055649147 0.0000000
## c009:3:-c006:5dot6:     -0.151019686 -0.189629072 -0.112410300 0.0000000
## c009:5dot6:-c006:5dot6: -0.074638784 -0.113248170 -0.036029398 0.0000070
## c008:5dot6:-c008:3:      0.066447238  0.027837852  0.105056624 0.0000607
```

```
## c009:3:-c008:3:           0.009686086 -0.028923300  0.048295472 0.9920496
## c009:5dot6:-c008:3:       0.086066987  0.047457601  0.124676373 0.0000003
## c009:3:-c008:5dot6:      -0.056761152 -0.095370538 -0.018151766 0.0007441
## c009:5dot6:-c008:5dot6:  0.019619749 -0.018989637  0.058229135 0.7331751
## c009:5dot6:-c009:3:       0.076380901  0.037771515  0.114990287 0.0000044
```

I look at the variance within each isolate:pH category:

```r
# To see the variance for each category:

# Calculate variance for each category
variances <- tapply(sorted_df_091224$mumax, sorted_df_091224$Category, var)

# Print the variances
print(variances)
```

```
##      c005:3:  c005:5dot6:      c006:3:  c006:5dot6:      c008:3:  c008:5dot6:
## 3.106504e-05 1.357657e-03 1.148256e-04 1.495214e-03 3.103780e-05 2.496044e-04
##      c009:3:  c009:5dot6:
## 2.671656e-05 1.953235e-04
```
